# eScreen: a deep learning framework for functionally decoding the regulatory genome at single-nucleotide resolution

**DOI:** 10.64898/2026.02.02.703403

**Authors:** Shijie Luo, Liquan Lin, Han Zhang, Yiding Yang, Yunxiang Liu, Yingli Han, Fei Chen, Ruipu Liang, Ximing Nian, Jinxian Dai, Run Chen, Hongli Lin, Zhijie Li, Yongying Peng, Teng Fei, Jialiang Huang

## Abstract

The human genome is densely populated with cis-regulatory elements (CREs), yet deciphering their functional regulatory syntax and combinational logic remains a fundamental challenge. Here, we integrate 379 genome-scale CRISPR screen experiments, encompassing 21 million perturbations across 23 cell types, to construct a compendium of 41,239 high-confidence functional CREs from 530,527 candidates. Leveraging this resource, we develop eScreen, a deep learning model built on the StripedHyena2 architecture to functionally decode the regulatory genome at single-nucleotide resolution. eScreen achieves three primary functions: (1) predicts genome-wide cell-type-specific CRE functional activity with high accuracy, outperforming existing models; (2) provides mechanistic interpretation of regulatory syntax at single-nucleotide resolution; (3) dissects the functional organization of enhancer clusters through *in silico* perturbation analysis. We perform multiple independent CRISPR knockout, CRISPR interference (CRISPRi), and base editing screens to validate these functions of eScreen both at scale and on individual cases. Furthermore, we provide an interactive web server (https://escreen.huanglabxmu.com/) for the community to access the integrated CRISPR screen resources and eScreen functions. Collectively, our work establishes a highly precise and convenient tool to decode the causal effects of the regulatory genome.

## Introduction

A hallmark of eukaryote genome is the pervasive presence of noncoding cis-regulatory elements (CREs), which play indispensable roles in orchestrating spatiotemporal gene expression programs[1]. Far beyond passive DNA fragments, these elements serve as dynamic regulatory hubs, responding to developmental, environmental, and pathological cues[2]. Over the past two decades, large-scale international consortia—most notably the Encyclopedia of DNA Elements (ENCODE) project—have systematically annotated regulatory elements across the human genome, generating millions of putative CREs[3, 4]. However, the mere presence of a CRE does not equate to functional activity. These elements operate in inherently combinatorial and context-dependent manners, with regulatory activity varying dramatically across cell types, developmental stages, and disease conditions[2, 5].

Advances in diverse high-throughput technologies have rapidly expanded the toolkit for identifying functional CREs[6]. Chromatin immunoprecipitation followed by sequencing (ChIP-seq) enables detection of histone modifications and transcription factor binding[7]; STARR-seq measures CRE activity using reporter constructs[8]; and CRISPR-based screening technologies allow direct perturbation of genomic loci to assess functional effects[9]. Among these, CRISPR screens are particularly powerful due to their site-specificity, programmability, and scalability, enabling functional interrogation of CREs within their native chromatin context[10]. Notable studies have demonstrated the utility of CRISPR screen in mapping CREs of *BCL11A*[11] to uncover therapeutic targets for fetal hemoglobin regulation, and in dissecting CREs linked to genome-wide association study (GWAS) loci[12]. However, the rapid accumulation of CRISPR screen data, which are generated under heterogeneous editing strategies, target designs and phenotypic readouts, poses great challenges for data harmonization, functional interpretation, and cross-study integration[13]. The absence of a unified annotation framework and a generalized functional CRE roadmap limits the integration of regulatory insights across different biological contexts[14].

In parallel, computational models have advanced our ability to predict CRE activity. Methods such as NVWACE[15] and DeepSEA[16] integrate chromatin accessibility, histone modifications, and 3D contact data to generate genome-wide catalogs of CREs. Massively parallel reporter assays (MPRA) enable high-throughput and customizable sequence libraries for direct functional testing, giving rise to a new generation of deep learning models such as Malinois[17], which aims to learn the regulatory grammar. In addition, several large-scale predictive models have been developed to infer epigenomic profiles directly from DNA sequence, including AlphaGenome[18] and Enformer[19]. However, these approaches have inherent limitations: (1) epigenome-based models or sequence-to-profile models are constrained by the resolution of current chromatin assays and cannot assess functional regulatory relationships, leading to elevated false positive rates[20]; (2) reporter-based methods, on the other hand, rely on exogenous plasmid constructs that fail to recapitulate the native genomic and topological context[21]. Consequently, neither strategy fully captures the causal and context-specific regulatory potential of CREs within the endogenous genome.

To address these challenges, we present eScreen, the first sequence-based deep learning model trained on a comprehensive compendium of functional CREs integrated from large-scale CRISPR screens. eScreen predicts genome-wide CRE functional activity, resolves regulatory syntax at single-nucleotide resolution, delineates cell-type-specific transcription factor regulatory programs, and quantifies core functional CRE and synergistic interactions within enhancer clusters. We rigorously validate eScreen through multiple types of perturbation screens and detailed CRE characterization. This study provides a unified and scalable platform for decoding the causal functions of the noncoding genome, offering novel insights into transcriptional regulation and empowering the functional interpretation of genetic variation in health and disease.

## Results

### A compendium of functional cis-regulatory elements (CREs) integrated from 379 CRISPR screens

To systematically elucidate the function of CREs across the human genome, we integrated a comprehensive dataset of 379 noncoding CRISPR screens. These experiments encompass over 21 million perturbations across 23 cell types, employ 5 types of CRISPR systems, and feature diverse experimental designs (genome-wide, CRE focused or tiling) with phenotypic readouts mainly converging on “proliferation-based cell fitness” and “Fluorescence-Activated Cell Sorting (FACS)-based target signal” (**Fig. 1a**). By intersecting these functional data with the ENCODE registry of candidate CREs, we mapped 530,527 candidate CREs. Applying MAGeCK analysis[22], we identified 41,239 (∼7.8%) high-confidence functional CREs that exhibited significant phenotypic effects upon perturbation in at least one screen (**Fig. 1a**, **Methods**). This functional CRE compendium spans diverse CRISPR systems, cell types, regulatory classifications, and associated genes, underscoring the breadth and utility of the resource (**Fig. 1b**, **Extended Data Fig. 1a**). As an illustrative example, in K562 cells, we identified 17,414 (∼4.7%) functional CREs, whose perturbation affected either proliferation or fluorescence signal (**Fig. 1c-d**, **Supplementary Table 2**). These elements were enriched near genes encoding key hematopoiesis-related transcriptional regulators, such as *MYB, GATA1,* and *MYC*[23–25] (**Extended Data Fig. 1b**). Of note, the proportion of functional CREs varied across the 13 K562 datasets, which likely reflects differences in experimental design (e.g., genome-wide vs. tiling) and target CRE selection criteria (**Fig. 1d**).

**Fig. 1.**
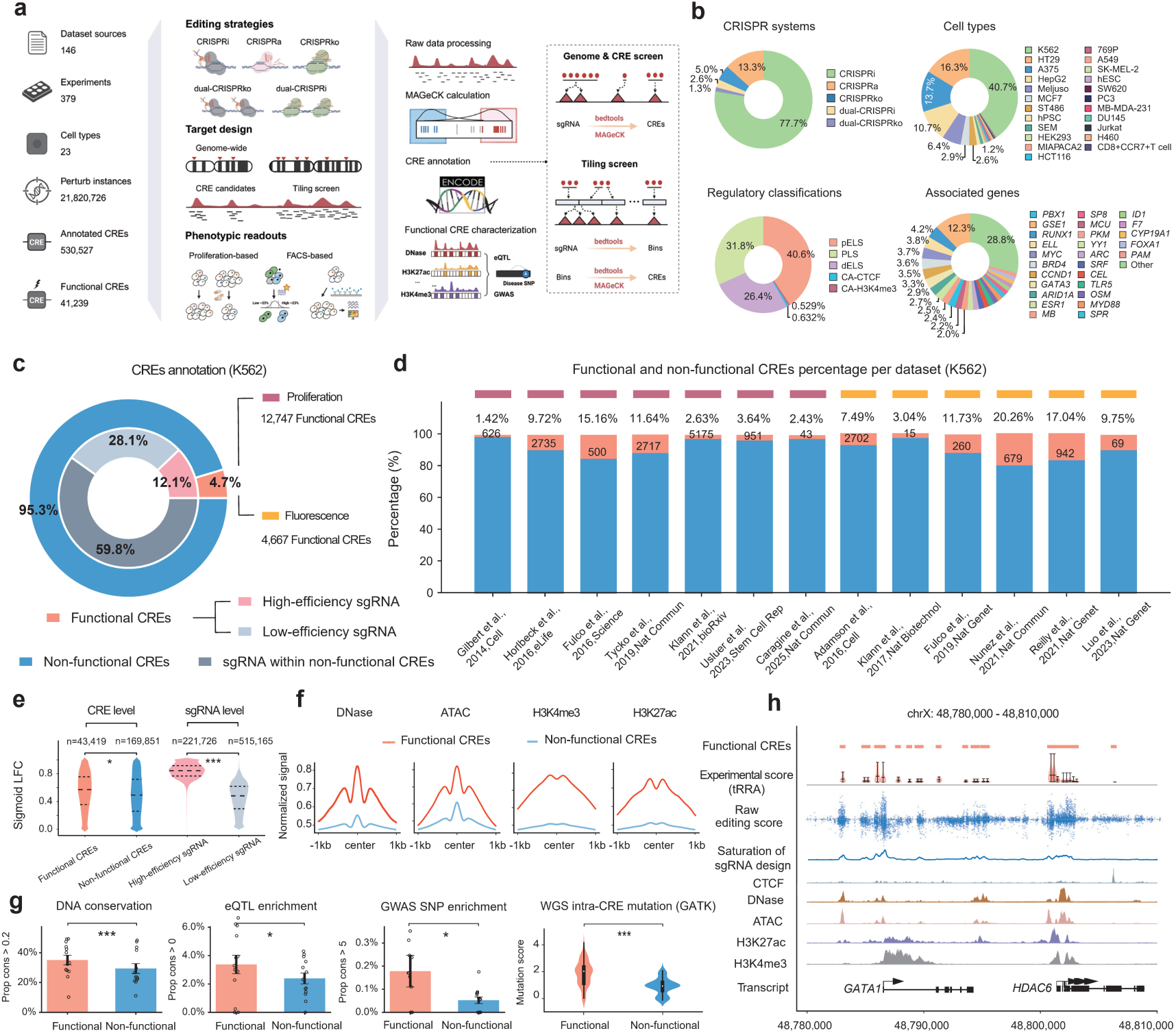
A compendium of functional cis-regulatory elements (CREs) derived from 379 CRISPR screens. **a**, Overview of the pipeline for defining functional CREs by integrating 379 noncoding CRISPR screens (>21 million perturbations across 23 cell types) with ENCODE CRE annotations. **b**, Summary statistics of the functional CRE compendium. Pie charts show the distribution of identified functional CREs grouped by CRISPR systems, cell types, regulatory classifications, and associated genes. **c**, Distribution of functional and non-functional CREs in K562 cells. sgRNAs targeting functional CREs were further categorized as high- or low-efficiency based on their perturbation effect. **d**, Functional CREs identified across 13 independent K562 datasets were grouped by their primary phenotypic readout (proliferation or fluorescence). The bar plot shows, for each dataset, the percentage of functional versus non-functional CREs. **e**, Editing efficiency of sgRNA targeting CREs. Violin plots compare editing efficiency (sigmoid LFC) between sgRNAs targeting functional and non-functional CREs, as well as between high- and low-efficiency sgRNAs. Statistical significance was assessed by t-test (\**p* < 0.05, \*\**p* < 0.01, \*\*\**p* < 0.001; *n.s.*, not significant). **f**, Enrichment of chromatin features at functional CREs in K562 cells. Normalized signal profiles for DNase-seq, ATAC-seq, H3K4me3, and H3K27ac ChIP-seq across ±1 kb centered on functional and non-functional CREs. **g**, Enrichment of functional genomic annotations in functional CREs. Functional CREs show significant enrichment for evolutionarily conserved regions, eQTLs, GWAS SNPs linked to hematopoietic traits, and recurrently mutated regions in acute myeloid leukemia (AML) from the GATK database. Each dot represents one K562 dataset. Statistical significance was assessed by t-test (\**p* < 0.05, \*\**p* < 0.01, \*\*\**p* < 0.001; *n.s.*, not significant). **h**, Functional CREs at the *GATA1* locus. Top, functional CREs (red bars) defined by integrating CRISPR screen compendium. Middle, a unified experimental score (transformed RRA score, tRRA), calculated by aggregating sgRNA activity (blue dots) across different K562 datasets and shown as red violin plots. Bottom, epigenomic tracks (CTCF, DNase-seq, ATAC-seq, H3K27ac and H3K4me3) are shown for context.

We next characterized functional CREs by associating their perturbation effects with essential genomic and epigenomic features. Using sigmoid-normalized log_2_ fold-change (sigmoid LFC) score, sgRNAs within functional CREs were categorized as high-/low-efficiency (**Fig. 1c**). As expected, sgRNAs in functional CREs showed significantly higher sigmoid LFC compared to sgRNAs in non-functional CREs (**Fig. 1e**). Within functional CREs, high-efficiency sgRNAs further exhibited higher scores than low-efficiency ones (**Fig. 1e**). Our functional CREs annotation was supported by their pronounced concordance with established epigenetic features, including chromatin accessibility and active histone modifications, relative to non-functional CREs (**Fig. 1f**). These functional CREs were also significantly enriched for evolutionarily conserved sequences, hematopoiesis-associated GWAS SNPs[26], eQTLs[27], and recurrently mutated regions in acute myeloid leukemia (AML) from the GATK database[28] (**Fig. 1g**, **Extended Data Fig. 1c**). For example, at the *GATA1* locus, we identified 23 functional CREs in K562 with high editing scores (blue dots) based on our integrated CRISPR screen compendium, in agreement with prior studies[9, 29–38] (**Fig. 1h**). Moreover, the experimental scores generated from our framework will serve as a gold standard benchmark for subsequent models of CRE activity (**Fig. 1h**). Collectively, these results provide a foundational resource for training predictive models and enabling the mechanistic interpretation of CRE function.

### eScreen, a deep learning model for predicting and dissecting functional CREs

Leveraging this compendium of noncoding perturbations, we developed eScreen, a StripedHyena2-based deep learning model[39] for genome-wide prediction of CRE functional activity from DNA sequences (**Fig. 2a-c**, **Methods**). StripedHyena2 employs a hierarchy of explicit and implicit convolutional operators to jointly capture short-, medium-, and long-range dependencies in sequence data, enabling scalable representation of distal regulatory interactions (**Fig. 2b**). Building on this foundation, eScreen integrates transcription factor motif embeddings as explicit regulatory priors with sequence-based modeling of multi-scale regulatory syntax (**Fig. 2a**). These signals are iteratively resolved through StripedHyena2 convolutional operations, allowing eScreen to capture cooperative motif interactions and long-range regulatory effects that shape CRE functional activity. Beyond sequence alone, eScreen can be augmented with an optional graph neural network (GNN) module to incorporate local chromatin context by integrating diverse epigenetic profiles (**Fig. 2a, b**, **Methods**). This extension explicitly models spatial relationships between CREs and proximal epigenomic features, enabling syntax-aware refinement of CRE activity predictions under native chromatin environments.

**Fig. 2.**
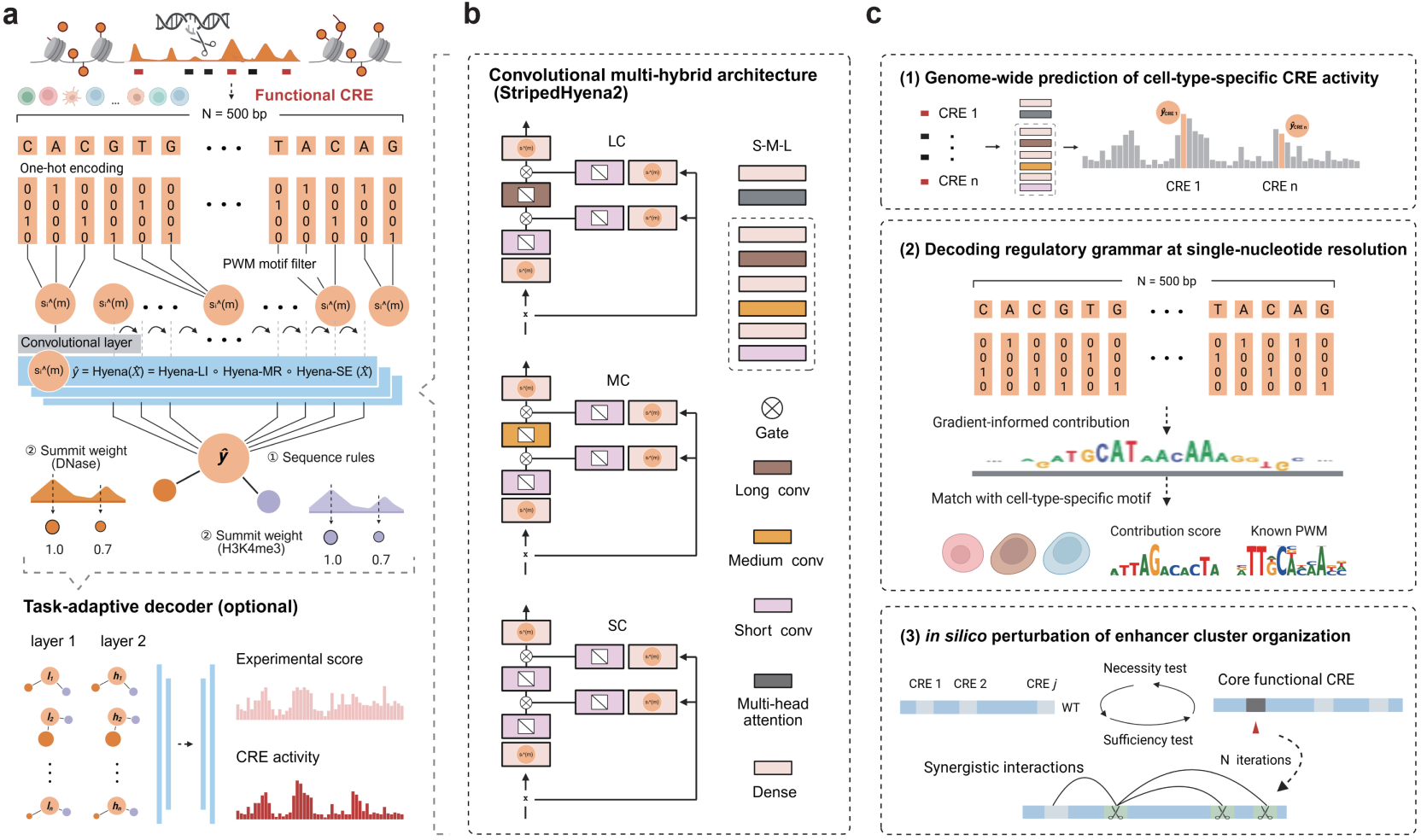
eScreen: a deep learning model for functionally decoding the regulatory genome at single-nucleotide resolution. **a**, eScreen training uses 500-bp sequences of functional and non-functional CREs identified in the CRISPR screen compendium, optionally augmented by a graph neural network that integrates epigenomic profiles to enhance prediction. **b**, eScreen is built on a StripedHyena2 convolutional architecture, which employs a hierarchy of explicit and implicit convolutional operators to capture short-, medium-, and long-range dependencies in sequence data and iteratively model cooperative motif interactions. **c**, The model performs three tasks: (**1**) predicting cell-type-specific CRE functional activity across the genome; (**2**) decoding regulatory grammar at single-nucleotide resolution via gradient-informed contribution analysis; and (**3**) dissecting the functional organization of enhancer clusters through *in silico* perturbation.

These components allow eScreen to extract high-resolution regulatory features while remaining scalable to genome-wide analyses. eScreen is formulated for three primary tasks (**Fig. 2c**): (1) predicting cell-type-specific CRE functional activity on genome-wide scale; (2) deciphering regulatory grammar at single-nucleotide resolution; and (3) dissecting the functional organization of enhancer clusters through *in silico* perturbation analysis. These complementary tasks enable both predictive and mechanistic interrogation of noncoding regulatory sequences within a unified framework. Distinct from previous methods[15–17, 19, 40], eScreen directly leverages noncoding CRISPR perturbation data to learn CRE regulatory syntax from native chromatin contexts, rather than relying on correlative epigenomic annotations or reporter assays. Together, by grounding sequence modeling in causal functional readouts, eScreen establishes a new paradigm for CRE modeling that bridges predictive accuracy with mechanistic interpretability.

### eScreen predicts cell-type-specific CRE activity

We benchmarked eScreen’s performance against existing state-of-the-art models for CRE functional activity prediction, including AlphaGenome[18], DeepSEA[16], Enformer[19], Malinois[17], NVWACE[15] and HyenaDNA[40]. In the sequence-only setting, eScreen outperformed these models, achieving superior accuracy (**Fig. 3a**). Performance was further improved by incorporating epigenetic features, resulting in robust predictive accuracy across heterogeneous cell types (**Extended Data Fig. 2a**). For example, eScreen reliably distinguishes active elements within the ∼2Mb *MYC* locus, despite the presence of numerous sequence-divergent enhancers (**Fig. 3b**). Such robust performance extends to several additional loci, including *EGR1*, *MNT*, *MXI1*, and *E2F1* (**Extended Data Fig. 2b**).

**Fig. 3.**
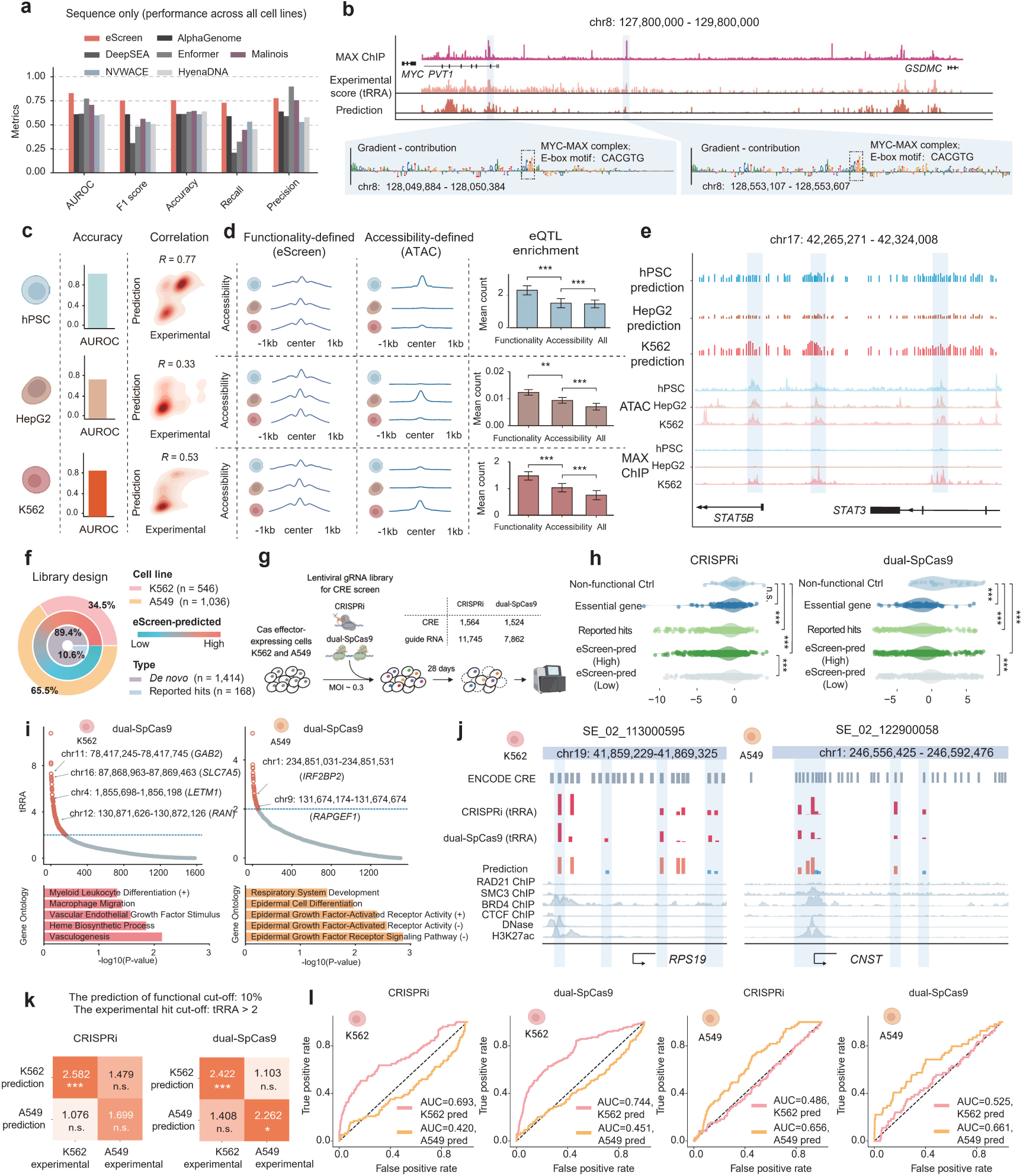
eScreen predicts cell-type-specific functional CREs from DNA sequence. **a**, Benchmarking of eScreen against existing tools, including AlphaGenome, DeepSEA, Enformer, Malinois, NVWACE and HyenaDNA for sequence-only prediction of CRE’s functional activity. All models were trained and evaluated on functional CRE sequences across all cell lines. **b**, eScreen-predicted functional activity across the ∼2Mb *MYC* locus, shown alongside MAX ChIP-seq profiles and CRISPR experimental score (tRRA). Gradient contribution analysis pinpointed nucleotide-level determinants predictive of MAX binding at two putative functional CREs, validated by independent MAX ChIP-seq data. **c**, Performance of eScreen in three representative cell types: K562, hPSC and HepG2. Density scatter plots compare eScreen-predicted scores (*y*-axis) with CRISPR experimental scores (*x*-axis). **d**, Comparison of two strategies for identifying cell-type-specific CREs: functionality-defined (by eScreen) vs. chromatin accessibility-defined (ATAC), with corresponding chromatin accessibility fitting curves and eQTL enrichment analyses. Significance was assessed using Mann-Whitney-U test (\**p* < 0.05, \*\**p* < 0.01, \*\*\**p* < 0.001; *n.*s., not significant). **e**, Predicted-regulatory landscapes at the *STAT3* locus, showing eScreen-predicted CRE activity, chromatin accessibility, and MAX ChIP-seq binding across three cell types. **f-g**, Schematic of the experimental workflow. (**f**) A total of 1,582 CREs within 205 super-enhancers in K562 and A549 cells (outer) were selected for functional testing using both CRISPRi and dual-SpCas9 editing systems, annotated with eScreen-predicted functional scores from low to high (middle), and 168 CREs (10.6%) were previously reported hits (**Supplementary Table 1**), whereas the remaining 1,414 CREs (89.4%) were *de novo* prediction by eScreen (inner). (**g**) Workflow for SEs screen in K562 and A549 cells. CRISPRi and dual-SpCas9 editing systems were used to target 1,564 and 1,524 CREs, respectively, employing a total of 11,745 sgRNAs and 7,862 pgRNAs. **h**, Distribution of log_2_ fold-change (LFC) values in CRISPRi and dual-SpCas9 screen systems in K562 cells for sgRNAs targeting non-functional control regions (Non-functional Ctrl), essential genes (Essential gene), previously reported functional CREs (Reported hits), and eScreen-predicted CREs (High/Low, 10% threshold). Statistical significance was calculated by t-test (\**p* < 0.05, \*\**p* < 0.01, \*\*\**p* < 0.001; *n.s.*, not significant). **i**, High-confidence functional CRE hits identified in K562 (left) and A549 (right) cells by dual-SpCas9 screening. RRA plots display transformed RRA score (tRRA) from MAGeCK analysis (Day 28 vs. Day 0). The labeled genes are adjacent to CRE hits detected by the ABC model. And bar plots show -log_10_ p-value derived from gene ontology (GO) enrichment analysis of all genes located adjacent to CRE hits. **j**, Two representative super-enhancer loci showing eScreen-predicted CRE functional activity, experimental score (tRRA) across CRISPRi and dual-SpCas9, and matched epigenetic profiles for context. **k**, Heatmap shows fold enrichment of experimental hits (tRRA > 2) from K562 and A549 cells among the top 10% of eScreen-predicted cell-type-specific functional CREs in CRISPRi (left) and dual-SpCas9 (right) editing systems. Significance was assessed using hypergeometric test (\**p* < 0.05, \*\**p* < 0.01, \*\*\**p* < 0.001; *n.*s., not significant). **l**, ROC curves evaluating eScreen’s predictions of cell-type-specific functional CREs, using experimental score identified from CRISPRi or dual-SpCas9 screen in K562 and A549 cells as the gold-standard.

We next evaluated eScreen across three representative cell types—HepG2 (human liver cancer cells), hPSC (human pluripotent stem cells), and K562 (human leukemia cells), in which eScreen consistently achieved high predictive accuracy (**Fig. 3c**). Interestingly, the cell-type-specific functional CREs predicted by eScreen (functionality-defined) displayed largely similar chromatin accessibility across different cell types, in contrast to CREs defined solely by chromatin accessibility (accessibility-defined) (**Fig. 3d**). Moreover, functionality-defined CREs showed higher enrichment for cell-type related eQTLs than accessibility-defined CREs (**Fig. 3d**), indicating that eScreen’s functionality-defined predictions better capture disease-relevant biology. For instance, at the *STAT3* locus, eScreen predicted three enhancers with highly functional activity specific to K562 cells, despite similar chromatin accessibility profiles across the three cell types. These predictions were corroborated by cell-type-specific MAX binding in K562 but not in hPSC or HepG2 cells (**Fig. 3e**). This suggested that eScreen captures subtle regulatory syntax and distinguishes cell-type-specific CRE activity by integrating CRISPR perturbation data.

### CRISPR screens validate the accuracy of eScreen on functional CRE prediction

To further validate the accuracy of eScreen’s prediction of cell-type-specific CRE activity, we sought to employ a systematic and unbiased experimental approach that relies on CRISPR-based perturbation screens. We selected 1,582 CREs spanning 205 super-enhancers (SEs) in either K562 or A549 (human lung cancer) cells, with eScreen predicted functional scores ranging from low to high (**Fig. 3f**). Notably, 1,414 (89.4%) of these CREs were previously underexplored in our integrated CRISPR screen compendium, providing a stringent test of eScreen’s *de novo* predictive capability (**Fig. 3f**). These candidate CREs were then targeted for loss-of-function perturbation in K562 and A549 cells using both CRISPRi and dual-SpCas9 (paired gRNA) screens, the currently optimal and complementary perturbation tools for deciphering CREs as determined in our benchmark study[41]. Functional CREs regulating the cell fitness in the two cell line models were identified (**Fig. 3f,g**, **Methods**, **Supplementary Table 3**). Both CRISPRi and dual-SpCas9 systems showed strong and concordant depletion for sgRNAs targeting essential genes (e.g., *RPS19*, *EEF2*) and previously reported top CRE hits, relative to non-functional control regions (*e.g.*, *AAVS1*, *ROSA26*), confirming the high data quality of our screens (**Fig. 3h**, **Extended Data Fig. 3a**). As expected, CREs with high eScreen-predicted scores exhibited significantly greater perturbation effect compared to those with low scores (**Fig. 3h**), indicating a strong overall agreement between prediction and experiment.

We next identified cell-type-specific CREs in K562 and A549 cells that likely regulate nearby genes essential for cellular fitness (**Fig. 3i** and **Extended Data Fig. 3b**). In K562 cells, CRE hits are enriched near genes involved in myeloid differentiation, macrophage migration, VEGF signaling, heme biosynthesis, and vasculogenesis, highlighting their role in hematopoietic and vascular function. Meanwhile, CRE hits in A549 cells regulate genes controlling respiratory development, epidermal differentiation, and EGF receptor signaling, underscoring their contribution to epithelial identity and growth factor dependent pathways (**Fig. 3i**). For example, at two SEs near the genes *RPS19* and *CNST*, we selected 8 and 9 CREs for validation. At these loci, the functional activities predicted by eScreen showed strong concordance with the experimental measurements (**Fig. 3j** and **Extended Data Fig. 3c**). Furthermore, eScreen-predicted cell-type-specific functional CREs were strongly enriched among the experimentally identified hits in their respective cell types (**Fig. 3k**). Examining the editing landscape for all CREs, eScreen’s predictions for K562-specific activity achieved the highest accuracy on K562 screening data (AUC = 0.693 for CRISPRi, 0.744 for dual-SpCas9), but showed lower accuracy for A549-specific predictions on the same data (**Fig. 3l**, **Supplementary Table 3**). Conversely, in A549 screens, A549-specific predictions performed best, whereas K562-specific predictions were less accurate, highlighting a clear cell-type-specific correspondence between predicted and observed CRE activity.

Collectively, these results demonstrate that eScreen robustly and accurately predicts functional CREs across cellular contexts, combining high generalizability with a strong capability to capture cell-type-specific regulatory logic.

### eScreen deciphers the regulatory grammar of CREs at single-nucleotide resolution

To decode the regulatory grammar of functional CREs, eScreen computes gradient-informed contribution scores within a 500-bp window for each element, achieving single-nucleotide resolution (**Fig. 4a**, **left**). We then scanned CRE sequences against 879 transcription factor (TF) position weight matrices (PWMs) from JASPAR and HOCOMOCO v12 databases[42, 43], linking sequence content to putative TF repertoires in each cell type (**Fig. 4a**, **middle**). Finally, cell-type-specific contribution profiles were factorized using Non-negative Matrix Factorization (NMF) to define higher-order regulatory programs that capture TF combinatorial logic (**Fig. 4a**, **right**).

**Fig. 4.**
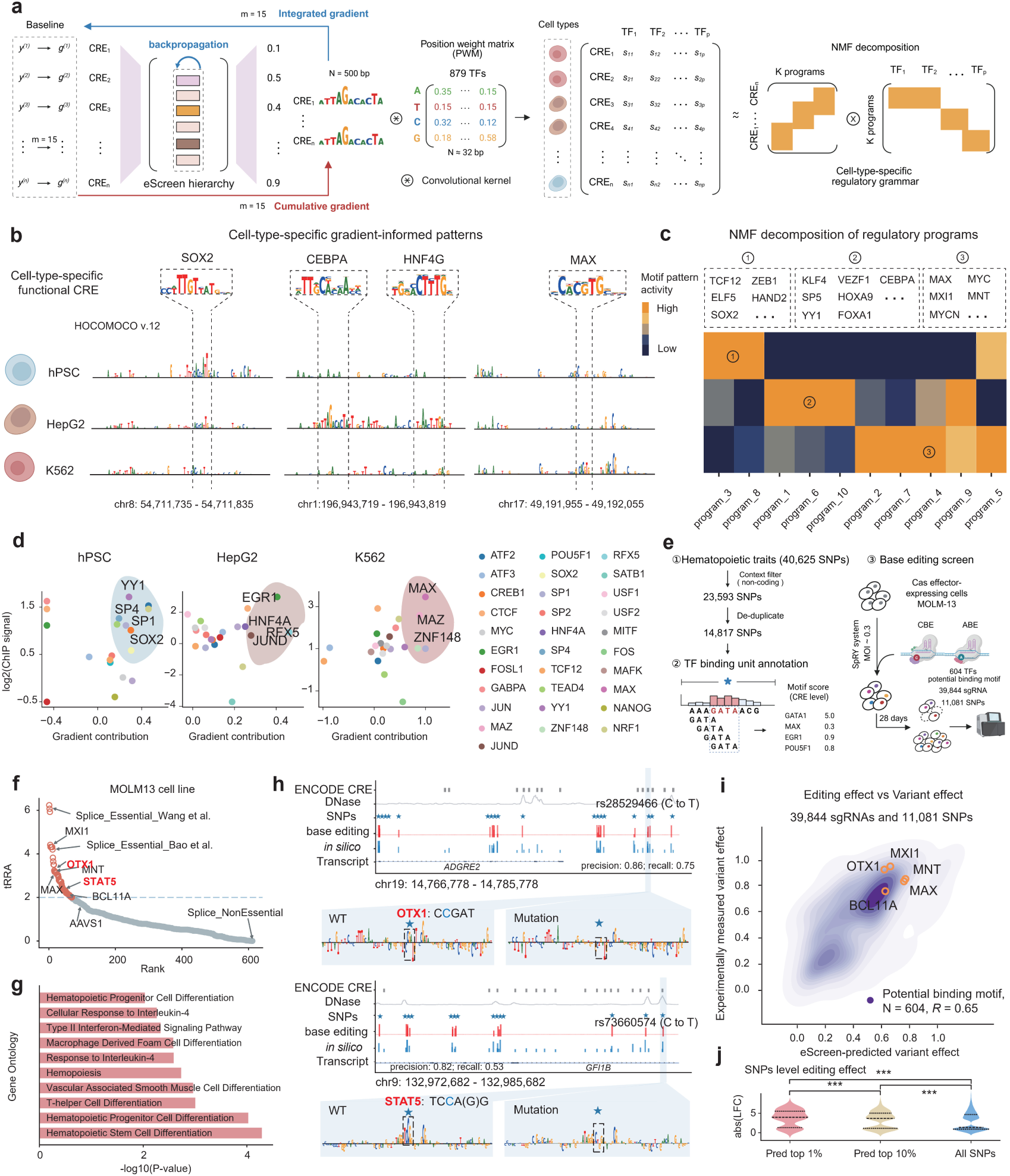
eScreen deciphers the regulatory grammar of CREs at single-nucleotide resolution. **a**, Schematic of the computational framework for decoding regulatory grammar using gradient-informed contribution analysis (**Methods**). **b**, Representative cell-type-specific gradient-informed patterns in K562, hPSC and HepG2 cells, with matched TF motifs from the HOCOMOCO v12 database. **c**, Non-negative matrix factorization (NMF) analysis identifying cell-type-specific motif combinatorial programs, with distinct cell-type-specific motifs indicated by dashed boxes. **d**, Validation of cell-type-specific motif predictions. Scatter plots show the eScreen-derived gradient contribution scores for predicted TF motifs (*x*-axis) with independent ChIP-seq binding signals (*y*-axis). **e**, Workflow for GWAS SNP selection and base editing screen in hematopoietic cells. Hematopoietic trait-associated SNPs (n = 40,625) were filtered to retain noncoding variants and deduplicated, yielding 14,817 SNPs. SNPs were annotated for TF binding motifs (e.g., GATA1 motif as shown). Candidate SNPs were targeted with cytosine (CBE) or adenine (ABE) base editors using two independent gRNA libraries, respectively, in MOLM-13 cells (MOI 0.3), and cell fitness was analyzed at Day 28 post-transduction. **f**, TF motifs ranked by tRRA (experimental score) from the base editing screen in MOLM-13 cells. Labeled motifs were previously validated in hematopoietic cells[11, 50, 53, 55, 56]. **g**, Bar plots illustrate the -log_10_ p-value resulting from gene ontology (GO) enrichment analysis of all TFs annotated to the SNPs. **h**, *In silico* nucleotide mutagenesis by eScreen was performed on CREs surrounding *ADGRE2* and *GFI1B* loci, showing the predicted changes in regulatory activity (blue bar), and the corresponding phenotypic effect measured in base editing experiments (red bar). For CREs lacking SNPs, multiple A- or C- containing sequences were mutated, and the mean effect was quantified (blue bar without SNP). For example, rs28529466 and rs73660574 illustrate base contribution changes via *in silico* nucleotide mutagenesis. **i**, Correlation between the regulatory impact predicted by eScreen’s *in silico* mutagenesis and observed proliferation changes in base editing screens at motif level. Each point represents a motif, showing eScreen-predicted variant effect (x-axis) and the proliferation effect measured experimentally (y-axis). Representative screen-identified motif hits are labelled. **j**, Distribution of observed base editing effects (absolute LFC, abs(LFC)) for the top 1% and 10% of SNPs ranked by eScreen-predicted variant effect, compared with all SNPs. Significance was assessed using Mann-Whitney-U test (\**p* < 0.05, \*\**p* < 0.01, \*\*\**p* < 0.001; *n.*s., not significant).

We applied this framework to three representative cell types: hPSC, HepG2, and K562. The gradient-informed contribution scores quantified the regulatory influence of individual TFs at single-nucleotide resolution, successfully recovering canonical lineage-defining regulators, including SOX2 in hPSC cells[44], CEBPA and HNF4G in HepG2 cells[45], and MAX dimeric motifs in K562 cells[23] (**Fig. 4b**). NMF analysis further identified cell-type-specific combinatorial regulatory motifs enriched within eScreen-predicted functional CREs. In hPSC, motifs for ZEB1[46] and ELF5[47] were prominent; HepG2 showed enrichment of CEBP and FOXA family motifs[45]; and K562 cells exhibited a broader repertoire of canonical co-regulatory motifs, notably including MAX and MYC[23] (**Fig. 4c**, **Supplementary Table 4**). The heterogeneous programs identified by NMF exhibited distinct co-regulatory grammars that closely corresponded to the functional context of the originating cell type, such as stemness and embryonic regulation in hPSC, liver metabolism in HepG2, and hematopoietic-related pathways in K562 (**Extended Data Fig. 4a**). Moreover, eScreen’s gradient-informed contributions preferentially recovered transcription factors with strong, cell-type-specific occupancy, exhibiting high concordance with publicly available ChIP-seq profiles (**Fig. 4d**, **Extended Data Fig. 4b**). Together, these results demonstrate that eScreen’s gradient-informed contribution analysis captures the TF regulatory syntax of CREs at single-nucleotide resolution.

### Base editing screens confirm eScreen’s prediction on disease-relevant noncoding variants

Previous studies indicate that the vast majority of single nucleotide polymorphisms (SNPs) from GWAS reside in noncoding regions[48], suggesting their regulatory roles in shaping phenotypes through mechanisms such as modulation of TF binding affinity[48, 49]. To experimentally validate the power of eScreen at the single-nucleotide resolution, we curated candidate SNPs that were associated with hematopoietic traits according to GWAS Catalog[26]. After restricting the candidates to noncoding variants and removing redundancies, we obtained 14,817 unique SNPs (**Fig. 4e**, **left**), with each one annotated for putative TF binding motifs. Using the base editing efficiency predictor (BEEP)[50], we designed two pooled libraries of 39,844 sgRNAs targeting to install 11,081 SNPs spanning 604 TF motifs for adenine (ABE)[51] and cytosine (CBE)[52] base editors (**Fig. 4e**, **right**). The libraries were lentivirally transduced into ABE- or CBE-expressing MOLM-13 (acute myeloid leukemia) cells at low multiplicity of infection (MOI of 0.3) to install the targeted SNPs, and their effects on cell fitness were determined at the end of the screens (**Fig. 4e**, **right**). In addition to positive controls targeting known essential sites[50, 53, 54], the screens identified 66 motif-level hits, including several well-known hematopoiesis-related regulators (e.g., MXI1, MNT, OTX1, BCL11A and STAT5)[11, 50, 53, 55, 56] (**Fig. 4f-g, Extended Data Fig. 4c, Supplementary Table 5**).

We next directly compared eScreen’s *in silico* mutagenesis predictions with the phenotypic effects of each allelic variant measured in the base editing screen. At the *ADGRE2* and *GFI1B* loci, eScreen predicted higher contribution scores for GWAS variants than for adjacent CRE mutations outside GWAS regions, and these predictions closely matched the observed phenotypic effects (*ADGRE2* locus: precision 0.86, recall 0.75; *GFI1B* locus: precision 0.82, recall 0.53) (**Fig. 4h**). For example, eScreen accurately captured two SNPs (rs28529466 and rs73660574) within OTX1 and STAT5 motifs, recapitulating the effects of single-nucleotide substitutions and reflecting marked changes in the motif landscape (**Fig. 4h**, **Extended Data Fig. 4d**). Notably, SNP rs73660574 and its surrounding sequence harbor a STAT5-recognizable motif, suggesting that this locus may serve as a STAT5 binding site. STAT5 integrates cytokine signals to regulate proliferation and differentiation[57], while GFI1B acts as a proto-oncogenic transcriptional repressor essential for erythroid and megakaryocytic lineage specification[58]; both factors contribute to the coordinated control of proliferation and lineage commitment in these progenitor cells[59]. We hypothesize that rs73660574 modulates STAT5 binding at the *GFI1B* locus, altering *GFI1B* expression or activity. Such mutagenesis could reshape the affinity of the co-regulatory complex, ultimately influencing leukemia cell proliferation and differentiation (**Fig. 4h**). Overall, eScreen robustly captured the functional impact of single-nucleotide mutations across the 604 motifs (*R* = 0.65, **Fig. 4i**). Across all 11,081 SNPs, variants with high eScreen-predicted gradient-informed contributions showed stronger functional effects upon base editing compared with those with low contributions (**Fig. 4j**). Collectively, these base editing screens validate that eScreen’s gradient-informed contribution analysis deciphers CRE regulatory grammar at single-nucleotide resolution, accurately pinpointing TF binding events and prioritizing disease-relevant noncoding variants.

### *In silico* perturbation by eScreen dissects the functional organization of enhancer clusters

Many enhancers exist as clusters in the genome to regulate cell identity and disease-associated genes, often termed as enhancer clusters or super-enhancers (SEs)[60–64], yet the internal regulatory architecture remains incompletely characterized. To dissect this complexity, we leveraged eScreen’s ability to model CRE regulatory grammar and adapted an *in silico* perturbation framework[65] to pinpoint core functional CREs within enhancer clusters or SEs. This strategy jointly considers two complementary analyses: a necessity test, in which individual CREs are sequentially masked to identify indispensable elements, and a sufficiency test, in which all other CREs are masked to assess each CRE’s autonomous regulatory potential (**Fig. 5a**). We define “core functional CREs” as eScreen-predicted functional CREs that are both necessary and sufficient for driving regulatory activity within the enhancer cluster. Applying this framework to 3,309 enhancer clusters in K562 cells, comprising 2,578 eScreen-predicted functional CREs and 26,004 non-functional CREs (**Methods**), we identified 653 core functional CREs (**Fig. 5b**). Perturbations of these core functional CREs produced markedly stronger phenotypic effects than perturbation of other CREs within the same enhancer clusters (**Fig. 5b**). This trend was consistent across enhancer clusters adjacent to key hematopoietic regulatory genes, with core functional CREs exerting substantially stronger effects than the rest of the CREs (**Extended Data Fig. 5a**). To illustrate this pattern at the level of individual SEs, we presented the *EPHA2*-associated SE and pinpointed three core functional CREs (E1–E3), whose functional importance in promoting tumor progression has been previously reported through *in vivo* CRISPR/Cas9-mediated deletion[66] (**Fig. 5c**).

**Fig. 5.**
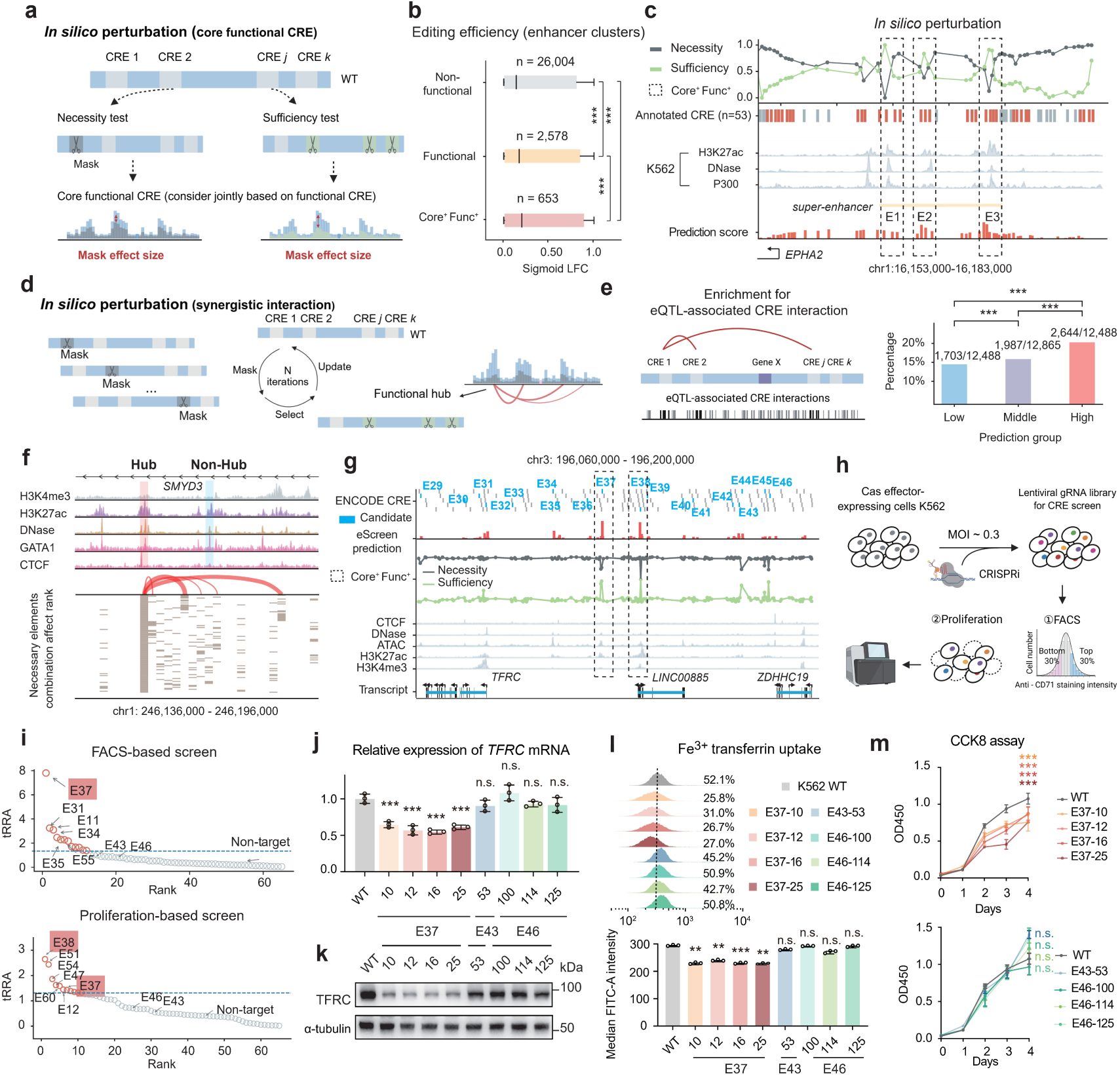
Dissecting functional organization of enhancer clusters by *in silico* perturbation. **a**, Schematic of the *in silico* masking strategy for pinpointing core functional CREs within enhancer clusters or super-enhancers, using necessity and sufficiency tests. **b**, Comparison of editing efficiency (sigmoid LFC) in integrated CRISPR screen dataset of sgRNAs targeting core functional CREs, functional CREs, and non-functional CREs within enhancer clusters in K562. Significance was assessed using Mann-Whitney-U test (\**p* < 0.05, \*\**p* < 0.01, \*\*\**p* < 0.001; *n.*s., not significant). **c**, *In silico* perturbation analysis at the *EPHA2* super-enhancer identifies core functional enhancers E1, E2, and E3, previously validated through *in vivo* CRISPR/Cas9-mediated deletion [66]. **d**, Schematic of the iterative *in silico* masking strategy for deconvolving synergistic enhancer interactions within an enhancer cluster. **e**, Enrichment of eQTL-inferred synergistic enhancer pairs among three enhancer combination stratified by eScreen-predicted interaction potential: low, middle, high. Significance was assessed using a chi-square test (\**p* < 0.05, \*\**p* < 0.01, \*\*\**p* < 0.001; *n.*s., not significant). **f**, *In silico* perturbation analysis at the *SMYD3* locus, revealing an enhancer (red bar) with extensive synergistic interactions, previously reported as a functional hub through *in vitro* CRISPR perturbation assays[68]. Supporting epigenetic tracks are shown. The heatmap ranks the top 100 enhancer combinations that most effectively restore regulatory activity; red lines indicate high-frequency co-occurrence edges, with the most recurrent CRE designated as the functional hub enhancer. **g**, *In silico* analysis of the *TFRC* locus using eScreen. Tracks display eScreen-predicted CRE functional activity, necessity, and sufficiency scores, nominating E37 and E38 as core functional enhancers. **h**, Experimental design for parallel CRISPRi screening strategy. A custom sgRNA library was used in FACS-based (monitoring *TFRC* coding protein CD71 surface expression) and proliferation-based screens. **i**, Identification of functional CREs. MAGeCK scatter plots highlight CRE hits from FACS-based (top, e.g., E37) and proliferation-based (bottom, e.g., E38 & E37) CRISPRi screens. **j-m**, Functional consequences of CRE knockout in single clonal K562 cells. (**j**) *TFRC* mRNA levels (RT-qPCR). (**k**) *TFRC* protein abundance (Western blot). (**l**) Cellular Fe^3+^ uptake. (**m**) Cell proliferation measured by CCK8 assay. For **j-m**, data were normalized to wild-type controls, and statistical significance was assessed using one-way ANOVA followed by Dunnett’s multiple comparisons test (\**p* < 0.05, \*\**p* < 0.01, \*\*\**p* < 0.001; *n.s.*, not significant).

Furthermore, eScreen can prioritize combinations of CREs with synergistic effects within enhancer clusters via iterative *in silico* perturbation (**Fig. 5d**). To validate these predicted synergies, we collected human blood eQTL data from the GTEx portal[27] and compiled eQTL-based synergistic enhancer pairs via a linear regression model following established methods[67] (**Fig. 5e**). We observed that eScreen-predicted high-potential interactions exhibited significant enrichment of eQTL-based synergistic enhancer pairs (**Fig. 5e**, **Extended Data Fig. 5b**, **Supplementary Table 6**). This capability is exemplified by the *SMYD3*-associated SE, in which we selected the top 100 CRE combinations showing the strongest restoration of activity and found that functional hubs were highly conserved within these combinations (**Fig. 5f**). This observation is consistent with our prior CRISPR/Cas9-mediated experiments, which identified that specific enhancer as a functional hub[68] (**Fig. 5f**). We further benchmarked eScreen against a recent multiplexed CRISPRi screen that quantitatively mapped enhancer-enhancer interactions across the *MYC* locus[69], and observed strong concordance between the eScreen’s predicted perturbation scores and experimental measurements, particularly for synergistic pairs such as E3/E4 and E7 (**Extended Data Fig. 5c**). Collectively, these findings establish eScreen’s *in silico* perturbation as a powerful approach for identifying core functional CREs and their cooperative interactions within enhancer clusters, providing a scalable framework for deciphering the logic of transcriptional regulation.

### eScreen identifies a core functional CRE in the *TFRC* enhancer cluster

To experimentally validate eScreen’s capacity to identify core functional CREs within complex enhancer architectures, we focused on the transferrin receptor (*TFRC*) gene locus, a pathogenic node in acute myeloid leukemia (AML)*. TFRC* is upregulated in AML, driving excessive iron uptake and elevated intracellular iron burden[70]. *In silico* perturbation analysis by eScreen resolved the *TFRC* enhancer cluster and highlighted E37 and E38 as the top functional CREs among 15 predicted functional CRE candidates (**Fig. 5g**). Next, we constructed a focused sgRNA library for targeted inhibition of 63 CREs within the *TFRC* enhancer cluster by a CRISPRi system (**Supplementary Table 7**). CRISPRi screens were performed in K562 cells with “proliferation-based cell fitness” and “FACS-based *TFRC* expression” as two functional readouts (**Fig. 5h**, **Extended Data Fig. 5d**). Overall, sgRNAs targeting predicted functional CREs exhibited significantly higher sigmoid LFC values than those targeting non-functional CREs, with core functional CREs showing the highest values (**Extended Data Fig. 5e**, **left**). And this pattern was also consistently validated in SE screening data shown in **Fig. 3g**, confirming the robustness of these core functional CRE identification (**Extended Data Fig. 5e**, **right**). Interestingly, E37 emerged as a uniquely recurrent and statistically significant hit across both screening modalities (the 1^st^ hit in FACS-based screen; 7^th^ hit in proliferation-based screen) (**Fig. 5i**), while E38 ranked the 1^st^ in fitness screen but not identified as functional in FACS screen (**Fig. 5i**). It is likely that E38 does not regulate *TFRC* expression as it resides obviously further to *TFRC* than to *LINC00885* (just within the promoter) (**Fig. 5g**). These results clearly confirm the functional importance of the two *in silico* predicted core functional CREs (E37 and E38) under corresponding scenarios. In contrast, elements such as E43 and E46, predicted by eScreen as non-functional, exhibited minimal effects across both screens (**Fig. 5i**).

To further validate these results, we performed targeted deletions of individual CREs by CRISPR knockout, and established single clonal K562 cell lines with genetic ablation of E37, E43 or E46 (**Extended Data Fig. 6a**, **Supplementary Table 7**). Compared with wild-type (WT) control cells, multiple lines of E37-deleted cells resulted in marked reduction in *TFRC* transcript abundance (**Fig. 5j**) and protein levels (**Fig. 5k**), whereas cells with perturbation of E43 or E46 all produced no measurable changes (**Fig. 5j, k**). Furthermore, loss of E37, but not E43 or E46, further impaired Fe³⁺uptake and significantly slowed cell proliferation (**Fig. 5l, m**), which are consistent with *TFRC*’s role as the principal iron importer and essential function in hematopoietic cells[71]. Indeed, shRNA-mediated knockdown of *TFRC* caused reduced Fe³⁺ uptake and cell proliferation (**Extended Data Fig. 6b**-**e**). These data corroborated that E37 is a core functional CRE via regulating *TFRC* transcription and expression.

Together with the preceding *in silico* analyses, these results validate eScreen’s ability to resolve core functional CREs within enhancer clusters and establish E37 as a critical regulatory node controlling *TFRC*-driven iron metabolism and proliferative capacity in hematologic malignancy.

### An eScreen web server for accessing and predicting functional CREs

To enable public access to our integrated CRISPR screening resources and eScreen predictions, we developed an interactive web server (https://escreen.huanglabxmu.com/). This server provides unified data management, prediction, and visualization of CRE function (**Fig. 6**). The platform aggregates diverse CRISPR screen datasets spanning multiple experimental designs, cell types, perturbation strategies, CRE annotations, and phenotypic readouts (**Fig. 6a**). It offers a structured search interface for efficient querying of gene- or CRE-centric information (**Fig. 6b**). In parallel, eScreen implements predictive models to infer cell-type-specific CRE activity, enabling users to explore genome-wide functional variability across cellular contexts (**Fig. 6c**). Predicted CRE activities are accompanied by integrated transcriptomic and epigenomic features, including histone modifications, TF binding, and chromatin accessibility, facilitating mechanistic interpretation of regulatory function (**Fig. 6d**). Together, the eScreen web server provides an end-to-end resource for systematic exploration of CRE function and regulatory architecture across human cell atlas.

**Fig. 6.**
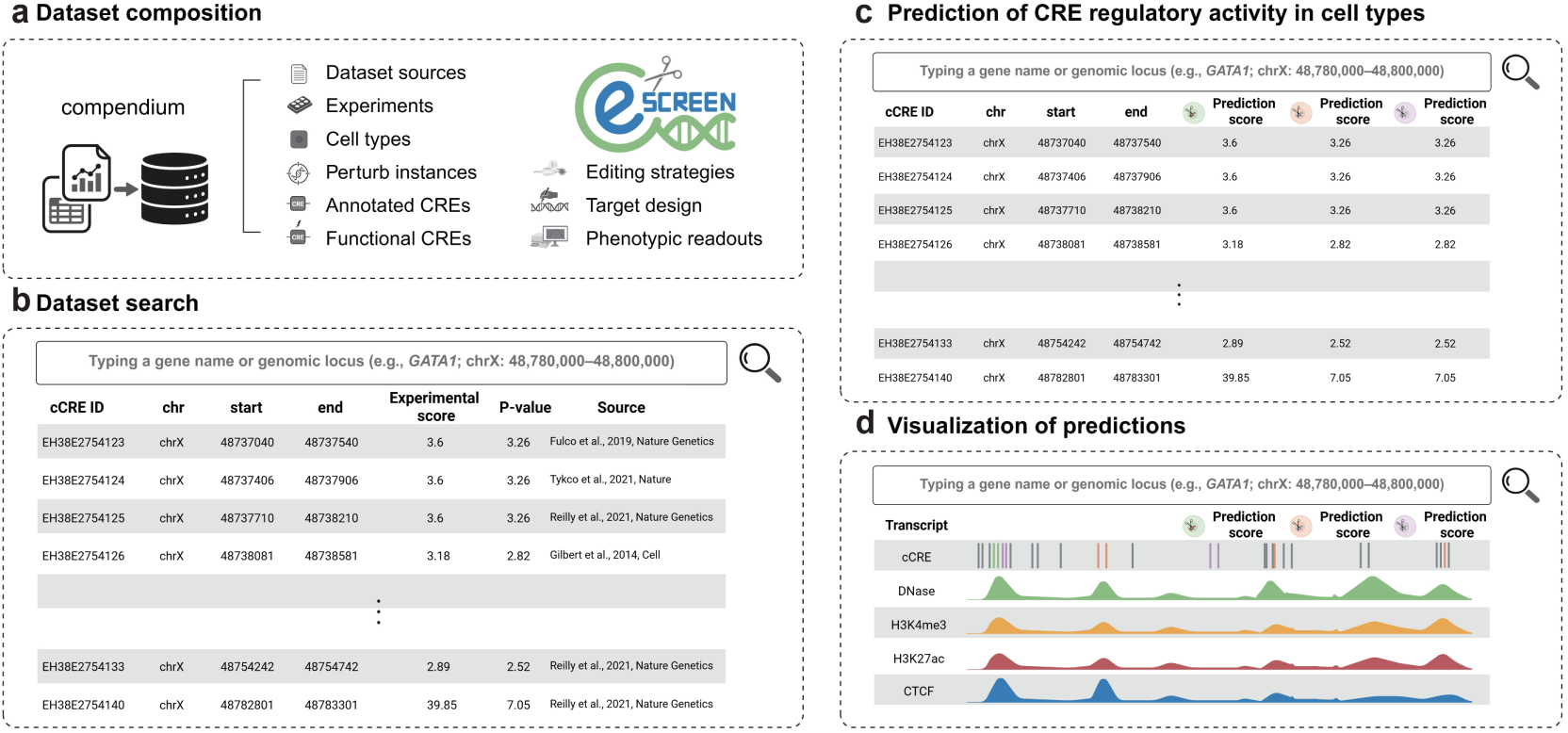
eScreen web server (https://escreen.huanglabxmu.com/): integrated CRE data management, prediction, and visualization. **a**, eScreen web server aggregates diverse datasets—including different experiments, cell types, perturbations, annotated and functional CREs, editing strategies, target designs, and phenotypic readouts. **b**, The platform provides a structured search interface for efficient gene- or CRE-centric data queries. **c**, eScreen predicts CRE activity across cell types and queries these predictions. **d**, Users can examine CRE-associated transcript and epigenomic features, such as histone modification, transcription factor binding, and chromatin accessibility, to interpret CRE function in specific cellular contexts.

## Discussion

Cis-regulatory elements (CREs) constitute the fundamental syntax of the noncoding genome, yet their functional resolution remains limited by cellular heterogeneity, chromatin context, and fragmented validation resources[2, 5]. Over the past two decades, large-scale consortia—most notably ENCODE—have generated millions of putative CREs[3, 4], but these annotations rely largely on correlative epigenomic signatures rather than direct perturbation-based evidence. A recent update from the ENCODE4 functional characterization centers conducted 108 screens in human cell lines[72]. Here, to our knowledge, we compiled the most comprehensive and coherent perturbation datasets for CREs, by integrating 379 CRISPR screens comprising >21 million perturbations. This resource provides a gold-standard functional foundation that overcomes previous heterogeneity and establishes a core reference for regulatory annotation.

Compared to reporter-based assays such as STARR-seq that quantify CRE activity outside its endogenous context[8], CRISPR-based technologies enable direct, programmable, and scalable perturbation of genomic loci within native chromatin[73]. Leveraging our comprehensive compendium, we introduce eScreen, a deep learning model trained explicitly on functional data from CRISPR screens to predict CRE activity directly from sequence. This framework transcends prior approaches that infer function indirectly from epigenomic surrogates[16] or artificial reporter assays[17].

The robustness of eScreen predictions is validated through high-throughput validation across multiple editing platforms, such as base editing[51], CRISPRi[34], and dual-SpCas9 knockout[41], on previously untested CREs. Beyond predictive accuracy, eScreen offers mechanistic interpretability. Gradient-based attribution reveals transcription factor binding syntax at single-nucleotide resolution. High-throughput base editing screens experimentally confirmed these fine-grained sequence determinants, demonstrating that the model captures causal regulatory grammar rather than correlative motifs. Coupled with *in silico* perturbation, eScreen systematically identifies core functional CREs and synergistic interactions within enhancer clusters, as exemplified at the *MYC* and *SMYD3* loci. This integrated framework enabled the discovery of E37 as a key regulator of iron metabolism and leukemic proliferation, linking a single noncoding element to a central disease phenotype. Moreover, eScreen provides a scalable and generalizable platform for prioritizing noncoding variants from GWAS[12], interpreting mutations of uncertain significance[74], and nominating therapeutic targets[75]. While eScreen provides a comprehensive and mechanistically interpretable framework for predicting CRE activity, it should be noted that the model is primarily trained on proliferation-based readouts, due to the constraint of currently available data sources. As a result, it may not fully capture the regulatory effects of CREs on the expression of specific target genes or their roles in other biological functions, such as differentiation, apoptosis, or stress responses. Future efforts integrating more available data on diverse phenotypic readouts, such as single-cell assays, lineage-specific screens, and *in vivo* models, will expand eScreen’s applicability and provide a more comprehensive map of CRE functionality[76]. Together, the compendium and predictive engine established here provide a unified and interpretable foundation for decoding the cis-regulatory code via bridging the genomic sequence with phenotypic output.

## Methods

### Compilation of functional cis-regulatory elements (CREs)

To construct a functional CRE compendium and leverage it for eScreen training, validation, and benchmarking, we compiled 379 CRISPR screen experiments from 146 published datasets (**Supplementary Table 1**). CRE coordinates were retrieved from the ENCODE candidate CRE registry to ensure consistent reference loci[3].

CRISPR libraries were categorized into genome-wide and tiling-based designs. For genome-wide screens, CRE activity was first computed based on original annotations, then mapped to CREs. For tiling-based screens, sgRNAs were first aligned to CREs, followed by element-level activity aggregation. All intersections were performed using bedtools intersect function[77] (**Fig. 1a**). The regulatory activity for each CRE was quantified using MAGeCK RRA[22] (default parameters).

To characterize functional CREs, we incorporated Phylogenetic conservation, eQTLs and GWAS data. Phylogenetic conservation was assessed using phastCons scores from the UCSC 100-way vertebrate alignment[78]. eQTLs data were obtained from the GTEx v8 release[27], and GWAS loci were downloaded from the NHGRI-EBI GWAS catalog[26].

### eScreen framework

We developed eScreen, a unified framework to define, predict, and characterize functional cis-regulatory elements (CREs) from large-scale CRISPR screen data. We assembled a compendium of noncoding CRISPR screens covering diverse editing strategies, target designs and phenotypic readouts, encompassing over twenty million editing events. The core subtasks of eScreen include: (1) predicting cell-type-specific CRE functional activity genome-wide; (2) deciphering CRE regulatory grammar at single-nucleotide resolution; and (3) dissecting functional organization of enhancer clusters through *in silico* perturbation (**Fig. 2c**).

### CRE activity prediction model

#### Model architecture

This prediction model consists of three functional components: (1) motif scanner, (2) StripedHyena2 module, and (3) task-adaptive decoder (**Fig. 2a-c**). Together, they enable accurate prediction and interpretation of functional CREs across diverse genomic and cellular contexts.

##### Motif scanner

To provide biologically informed input embeddings, we computed transcription factor (TF) binding scores for each nucleotide in the CRE sequence using a fixed-filter convolutional layer. Specifically, we implemented Fast Fourier Convolution (FFC) layer[79] with static filters initialized from position weight matrices (PWMs) of TF motifs curated from Vierstra, J., et al.[80]. Given a DNA sequence one-hot encoded a *X* ∈ ℝ^*L*×4^, where *L* = 500 bp, and PWMs motif filter *M_m_* ∈ ℝ*^k^*^×4^, the binding score at position at *i* for *m* motif *m* is computed via:

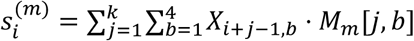

where *k* is the motif length, and *b* ∈ {*A*, *C*, *G*, *T*}. This convolutional scan generates a motif activation map *S_m_* ∈ ℝ^*L*-4+1^. To obtain position-wide embeddings, we further applied a max pooling operation over sliding windows to retain the strongest motif signals:

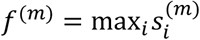

##### StripedHyena2

To capture long-range dependencies, we employed StripedHyena2[39], which models input-conditioned convolutions via Toeplitz matrix operations. The motif embeddings *f* are projected and concatenated to the sequence encoding*x*, serving as a biologically guided prior. The resulting tensor 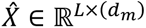 is passed into the StripedHyena2 module to model long-range dependencies among embedded motif signals and surrounding sequences. The Hyena modules operate on the motif-enriched sequence via:

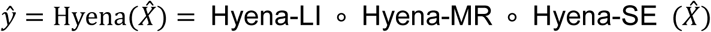

Each Hyena submodule is defined via a composition of input-conditioned convolutions:

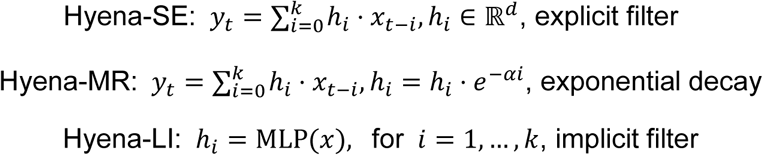

Finally, the multi-head attention layer integrates all contextual dependencies before decoding:

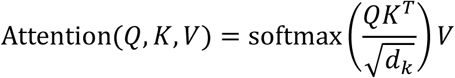

This model effectively combines motif-level priors, localized sequence features, and long-range regulatory dependencies, enabling robust activity prediction across varied CRE contexts.

##### Task-adaptive decoder

To accommodate task-specific flexibility, we implemented distinct decoding strategies. For multi-cell-line modeling, we employed a stack of fully connected linear layers to preserve generalizability across cellular contexts. For epigenomic integration tasks, we adopted a bipartite graph neural network (GNN)[81] that jointly models regulatory elements and epigenomic peaks through distance-constrained graph construction. Each CRE and epigenomic peak was represented as a node in graph *G* = (*V*, *E*), with nodes *V* = {*r_i_*}∪{*p_j_*}, where *r_i_* are regulatory elements and *p_j_* are epigenomic peaks. An edge *e_r,p_*∈*E* exists if:

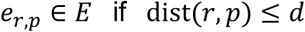

##### Model fitting

CRE sequences were padded to 500 bp on both sides and one-hot encoded. Epigenomic peak data were obtained from the ENCODE Project[82]. We used 80% of the data for training and 20% for testing. All models were trained using a single NVIDIA A30 24GB GPU.

##### Performance evaluation

The classification performance of eScreen was assessed using standard metrics, including area under the receiver operating characteristic curve (AUROC), precision, recall, f1 score, and accuracy. These metrics quantified both the model’s discriminative power and robustness, guiding iterative refinement of data preprocessing, sampling strategy, and hyperparameters.

### Gradient-informed contribution analysis

#### Motif contribution

To identify transcription factor (TF) motifs enriched within cell-type-specific CREs, nucleotide-level contribution scores were computed at the latent layer of StripedHyena2 using Integrated Gradients (IG) implemented via the Captum library[83]. IG quantifies the contribution of each input nucleotide by integrating gradients along a linear path from a baseline to the observed sequence, generating a 500 × 4 contribution matrix per CRE. Each sequence was scanned with JASPAR[42] core motifs to obtain per-base motif occurrence probabilities. For each motif, a weighted score was derived by aggregating base-wide contributions across the CRE. Scores were then averaged across CREs within each cell type to produce a cell-type-by-motif matrix.

#### Motif co-occurrence

To explore motif co-occurrence patterns, we constructed a motif co-occurrence matrix quantifying the frequency and contribution correlation of motifs appearing together within CREs. This analysis enabled the identification of putatively cooperative TF modules that may underlie combinatorial regulation.

#### Motif program

Non-negative matrix factorization (NMF)[84] was applied to the cell-type by motif matrix to uncover latent motif programs. Given a non-negative matrix *V* ∈ ℝ*^m^*^×*n*^ representing motif scores across *m* cell types and *n* motifs, NMF decomposes *V* into two non-negative matrices, *W* ∈ ℝ*^m^*^×*k*^ and *H* ∈ ℝ*^k^*^×*n*^, such that *V* ≈ *WH* where*k* denotes the number of motif programs. The factorization minimizes the Frobenius 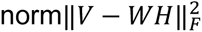, revealing motif combinations (rows of *H*) and their cell-type-specific activities (columns of *W*). Resulting motif programs were functionally characterized by Gene Ontology[85] enrichment analyses.

The above analysis pipeline was applied to three distinct cell lines: K562 (human leukemia cells), HepG2 (human liver cancer cells), and hPSC (human pluripotent stem cells), to validate the generalizability of the method and the robustness of cell-type-specific regulatory features (**Supplementary Table 4**).

### *In silico* perturbation

To evaluate the contextual importance of core functional CREs within an enhancer cluster, we conducted an *in silico* perturbation framework following the approach of Toneyan et al[65].

#### Necessity perturbation

Each element *r_i_* was masked and model predictions were recalculated to assess its contribution. Let *A* be the vector of predicted activities from the full sequence and 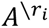 the prediction after masking *r_i_*. The necessity score was defined as the Pearson correlation:

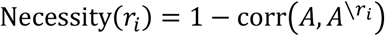

#### Sufficiency perturbation

All elements except *r_i_* were masked to assess its standalone contribution. Let 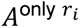 be the prediction when only *r_i_* remains:

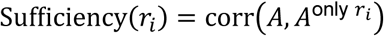

These scores reflect the context-dependent impact of each CRE in the model.

#### Synergistic interaction perturbation

Previous studies have shown that the activity of cis-regulatory elements (CREs) is approximately additive on average, such that the total regulatory activity of a CRE set can be well approximated by the sum of the activities of individual elements, with largely independent contributions[86, 87]. Under this additive assumption, perturbing a subset of CREs is expected to induce a proportional and relatively uniform change in the overall activity of the CRE set. In contrast, if a specific subset of CREs exhibits synergistic interactions, removing this subset would result in a disproportionately large change in the total predicted activity, exceeding the expectation derived from independent or additive effects. We leverage this deviation from additivity to quantify synergistic interactions through perturbation analysis. Formally, for a CRE subset of interest *E*, let **A** denote the vector of predicted activities prior to perturbation and **A’** the vector of predicted activities following the removal of *E*. We define the interaction score as the absolute difference between the summed predicted activities before and after perturbation:

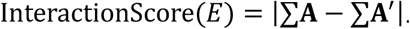

A higher interaction score indicates a stronger departure from additivity and suggests the presence of synergistic regulatory interactions within the perturbed CRE subset.

#### Enhancer cluster definition

Enhancer clusters, defined as aggregations of multiple enhancers forming three-dimensional chromatin domains[88], exhibit stronger cooperative effects in regulating their target genes than randomly assembled sets of enhancers, making them amenable to *in silico* perturbation analyses (**Supplementary Table 6**). Previously, enhancer clusters have been systematically identified across the human genome, with representative resources including dbSUPER[89], SEdb[90] and SEA[91]. Coordinates for enhancer clusters in the K562 cell line were retrieved from these three databases and overlapping regions were merged using the bedtools merge function. Enhancer clusters shorter than 30 kb were retained, resulting in a final set of 3,309 clusters for downstream *in silico* perturbation analyses.

### Benchmarking of eScreen against existing tools

All comparative analyses were conducted using the same input sequences as eScreen to ensure a fair and consistent evaluation across methods. We implemented the existing tools with their default setting following minimal data formatting adaptation required only for our experimental framework.

#### AlphaGenome[18]

CREs in the test set were mapped to their nearest transcription start sites (TSS). Masking-based evaluation was performed using AlphaGenome on the relevant output heads (e.g., gene expression and transcription initiation) to predict CRE regulatory activity across multiple functional genomic modalities. Since AlphaGenome does not currently offer direct predictions of CRE functional activity, we employed a masking-based strategy analogous to that used in Enformer[19].

#### DeepSEA[16]

Input sequences were modified to 512 bp DNA fragments, and multiple output heads were used to generate predictions for each cell line. As the model failed to converge with the default learning rate of 0.1, we trained it for 20 epochs using a learning rate of 0.00005, consistent with the eScreen training protocol.

#### Enformer[19]

CREs in the test set were mapped to their nearest transcription start sites (TSS). Gradient-based attribution was then performed using Enformer’s gradient attribution method on the CAGE output head corresponding to each cell line to predict CRE regulatory activity.

#### Malinois[17]

Input sequences were adjusted to 500 bp, and separate models were retrained for each cell line. Training was conducted using the default hyperparameters, with a learning rate of 0.00327 for 5 epochs.

#### NVWACE[15]

Input sequences were set to 500 bp, and multiple output heads were used to produce predictions for each cell line. Models were trained with default parameters, employing a learning rate of 0.001 for 20 epochs.

#### HyenaDNA[40]

The tiny-1k pretrained HyenaDNA model was fine-tuned, and, following the eScreen protocol, embeddings were generated via the embedding layer to distinguish outputs across cell lines. Fine-tuning was performed with a learning rate of 0.0001 for 20 epochs.

### Sigmoid-based normalization of gRNA activity

To enable quantitative comparison of gRNA performance across different readouts and selection directions, LFC values were further normalized using a sigmoid function, mapping the scores to a 0–1 scale. The transformation was defined as:

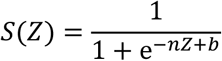

where *Z* represents the LFC for each sgRNA, and *n* and *b* are empirically determined parameters. These parameters were determined based on the global distribution of LFC values across different readouts and selection directions.

### Cell culture

K562 cells were cultured in RPMI-1640 medium supplemented with 10% fetal bovine serum (FBS), 1% penicillin-streptomycin, and 2 mM L-alanyl-L-glutamine. HEK293T and A549 cells were maintained in DMEM containing 10% FBS and 1% penicillin-streptomycin. Human AML cell lines MOLM-13 were cultured in RPMI-1640 medium supplemented with 10% FBS, 1% penicillin-streptomycin, and 2 mM L-alanyl-L-glutamine. All cells were incubated at 37°C in a humidified atmosphere with 5% CO₂. All cell lines were routinely tested and confirmed to be mycoplasma-free.

### CRISPRi and dual-SpCas9 screens for functional CREs

#### Plasmids and cell lines

Cas9 was expressed using the Lenti-Cas9-Blast vector (Addgene, #52962). The CRISPRi effector was delivered via a lentiviral construct encoding dCas9–KRAB–MeCP2, which was generated by inserting the dCas9m4-KRAB-MECP2 cassette into the Lenti-Cas9-Blast backbone following digestion with XbaI (Takara, #1634) and BamHI (Takara, #1605). A549 and K562 cell lines stably expressing either Cas9 or the CRISPRi effector were established by lentiviral transduction and Blasticidin (6 μg/mL; Solarbio, #B9300) selection for subsequent screening experiments.

#### Library design and construction

All sgRNAs and pgRNAs libraries were constructed using plentiGuide-BFP-Puro, which was derived from lentiGuide-Puro (Addgene #52963) by inserting the BFP sequence from pBA904 (Addgene #122238) at the SmaI and MluI restriction sites. All gRNAs were designed based on the human reference genome (GRCh38). To prioritize functional relevance and editing efficiency, sgRNAs were scored using CHOPCHOP[92] and DeepSpCas9[93], and pgRNAs were evaluated using DeepDC[41]. The final CRISPRi sgRNA library comprises 7,769 sgRNAs targeting 1,027 CREs in A549 and 3,976 sgRNAs targeting 537 CREs in K562. While the dual-SpCas9 pgRNA library contains 4,767 pgRNAs targeting 984 CREs in A549 and 3,095 pgRNAs targeting 540 CREs in K562 (**Supplementary Table 3**).

##### dual-SpCas9 library

Construction of the dual-SpCas9 paired guide RNA (pgRNA) library was performed using a two-step cloning strategy. Step I: the plentiGuide-BFP-Puro was digested with Esp3I (BsmBI; Thermo Fisher Scientific, #FD0454) at 37 °C for 1.5 h. The synthesized oligonucleotide pool was PCR-amplified and purified using the GeneJET PCR Purification Kit (Thermo Fisher Scientific, #K0702), followed by digestion with BpiI (BbsI; Thermo Fisher Scientific, #FD1014) at 37 °C for 1.5 h. Both vector and insert fragments were gel-purified using the GeneJET Gel Extraction Kit (Thermo Fisher Scientific, #K0692) and ligated with T4 DNA ligase (Thermo Fisher Scientific, #EL0016) at a 3:1 insert-to-vector molar ratio. The ligation products were introduced into self-prepared electrocompetent Stable *E. coli* cells by electroporation, achieving a library complexity corresponding to at least 50-fold coverage per designed construct. Transformed bacteria were expanded in ampicillin-containing LB medium at 30 °C for 16 h, and plasmid DNA was isolated using an Endo-Free Maxi Prep kit (TIANGEN, #4992194). Step II: plasmid libraries from Step I were subjected to a secondary Esp3I digestion at 37 °C for 1.5 h. Linearized and dephosphorylated vectors were resolved by 1.0% agarose gel electrophoresis and purified. A synthesized fragment containing the H1 promoter and gRNA scaffold was digested with Esp3I and ligated into the processed vector using T4 DNA ligase. Following bacterial transformation and plasmid purification, the dual-SpCas9 pgRNA library was obtained.

##### CRISPRi library

For construction of the CRISPRi sgRNA library, the plentiGuide-BFP-Puro vector was linearized and dephosphorylated, then assembled with PCR-amplified sgRNA oligonucleotides using 2× Gibson Assembly Master Mix at 50 °C for 1 h. Subsequent bacterial transformation, culture expansion, and plasmid purification were performed using the same procedures as described above.

#### Lentiviral transduction and high-throughput screening

For A549 screen, approximately 1.2 × 10⁷ A549 cells stably expressing Cas9 or the CRISPRi effector were transduced with lentiviral sgRNA or pgRNA libraries at a low multiplicity of infection (MOI ∼ 0.3), ensuring predominantly single-copy integration. Cells were selected with puromycin (1 μg/mL; Solarbio, #P8230) starting 48 h post-infection for 3 days, followed by a 2-day recovery period. A minimum of 6 × 10⁶ cells (corresponding to >500-fold library coverage) were harvested as the Day 0 reference population. The remaining cells were cultured for an additional four weeks without further selection to assess cell fitness during the screening period. The gRNAs of Day 0 and end point sample were PCR amplified and sequenced on an Illumina NovaSeq platform with PE150 mode.

For K562 screen, approximately 1.2 × 10⁷ K562 cells stably expressing Cas9 or the CRISPRi effector were transduced with lentiviral sgRNA or pgRNA libraries at a low multiplicity of infection (MOI ∼ 0.3) in the presence of polybrene (8 μg/mL; Yuanyebio, #R41131) to enhance transduction efficiency and ensure predominantly single-copy integration. Puromycin selection (1 μg/mL; Solarbio, #P8230) was initiated 48 h post-transduction and maintained for 3 days, followed by a 2-day recovery period. At least 6 × 10⁶ cells, corresponding to >500-fold library coverage, were harvested as the Day 0 reference population. The remaining cells were cultured for an additional four weeks without further selection to assess cell fitness throughout the screening period. The gRNAs of Day 0 and end point sample were PCR amplified and sequenced on an Illumina NovaSeq platform with PE150 mode. Two biological replicates were performed for these screens.

#### Data processing

Sequencing data generated from CRISPRi screens (FASTQ format) were processed using MAGeCK to generate raw sgRNA read count tables. For raw sequencing data from dual-SpCas9 screens, pgRNA sequences were first aligned using bowtie[94], followed by the removal of recombination reads. Raw log fold change (LFC) values for each sgRNA or pgRNA were calculated using MAGeCK. Subsequently, read counts were normalized and ranked using the robust rank aggregation (RRA) algorithm. Log10-transformed RRA scores (tRRA) were calculated to quantify the statistical significance of each target in promoting or suppressing the measured phenotype (**Supplementary Table 3**).

### Base editing screen

#### sgRNA design and library construction

We designed two independent base editing gRNA libraries for ABE and CBE to install targeted single nucleotide variants in the genome. These include variants associated with hematopoietic traits identified in genome-wide association studies (GWAS). We employed the SpRY system, which exhibits minimal PAM sequence constraints, thereby enabling sgRNA design without PAM restrictions at each target locus. For each candidate SNP, we generated up to 6 sgRNAs such that the target nucleotide fell within position 3-8 of the protospacer, the window of good editing efficiency for both ABE and CBE editors. All sgRNAs were designed against the human reference genome (GRCh38). To prioritize functional relevance and editing efficiency, we scored each sgRNA using BEEP (Base Editing Efficiency Predictor)[50], a deep learning-based model trained to predict base editing outcomes. Only sgRNAs with high BEEP scores were retained for library construction.

The final ABE library comprises 19,542 sgRNAs targeting 5,143 SNPs, while the CBE library contains 20,302 sgRNAs targeting 5,938 SNPs, these libraries include annotations for 604 potential transcription factor binding motifs (**Supplementary Table 5**).

For construction of the BE sgRNA library, the plentiGuide-BFP-Puro vector was linearized by BsmBI (New England Biolabs, #R0739L), dephosphorylated, and then assembled with PCR-amplified sgRNA oligonucleotides using 2× Gibson Assembly Master Mix (New England Biolabs, #E2611L) at 50 °C for 1 h. After incubation, the mix was transformed into self-prepared electrocompetent Stable *E. coli* cells (New England Biolabs, #C3040) by electro-transformation to reach the efficiency with at least 100X coverage representation of each clone in the designed library. The transformed bacteria were then cultured in LB medium with Ampicillin for 12∼14 h at 37°C. The library plasmids were then extracted with Endo-Free Maxi-prep Plasmid Kit (TIANGEN, #4992194).

#### Generation of base editor-expressing cells

Lentiviral particles encapsulating SpRY-ABE8e[50] or SpRY-evoA-BE4max[50] were produced in HEK293T cells by polyethylenimine (PEI)-mediated transfection. Viral supernatants were collected 48 h post-transfection, clarified by centrifugation, filtered, aliquoted, and stored at −80 °C. To generate stable base editor–expressing cell lines, MOLM-13 cells were infected with the corresponding lentiviruses at a low MOI (∼0.1) in the presence of polybrene, followed by blasticidin selection for 5–7 days. This yielded polyclonal base editor-expressing cell lines, which were subsequently expanded and used for high-throughput base editing screens.

#### High-throughput base editing screening

For functional screens, each base editor–expressing cell line was transduced with the corresponding sgRNA library: ABE-expressing cells with the ABE library and CBE-expressing cells with the CBE library. For each condition, 1.2 × 10⁷ cells were transduced at a low MOI (∼0.3) in the presence of 10 μg/mL polybrene, ensuring predominantly single-sgRNA integration while maintaining library representation. At 48 h post-transduction, cells were selected with 2 μg/mL puromycin for 3 days, followed by culture in fresh medium for 2 additional days. Selected cells were then maintained in puromycin-free complete medium for up to 28 days to allow base editing and downstream cell fitness effects to manifest. Genomic DNA was isolated from ∼6 × 10⁶ cells per sample, providing ∼300× library coverage. Integrated sgRNA cassettes were PCR-amplified with primers containing Illumina adapters and sample barcodes, pooled, and sequenced (paired-end 150 bp) on an Illumina NovaSeq platform. Sequencing reads were mapped to the reference sgRNA library to quantify guide abundance, and sgRNA representation at the endpoint (Day 28) was compared with the Day 0 input library. All screens were performed in two independent biological replicates.

#### Data processing

Sequencing data generated from base editing screens (FASTQ format) were processed using MAGeCK to generate raw sgRNA read count tables. Raw log fold change (LFC) values for each sgRNA were calculated using MAGeCK. To identify functional motifs, sgRNAs targeting the same motif were aggregated using the Robust Rank Aggregation (RRA) algorithm implemented in MAGeCK. Log10-transformed RRA scores (tRRA) were calculated to quantify the statistical significance of each variant in promoting or suppressing the measured phenotype (**Supplementary Table 5**).

### CRISPRi screening with *TFRC*-targeted sgRNA library

#### sgRNA design and library construction

A tiling CRISPRi library including 575 sgRNAs targeting the *TFRC* gene adjacent CREs (chr3: 196,054,000-196,214,000, hg38) was designed using CHOPCHOP[92], with all sgRNAs containing an NGG PAM sequence. A total of 58 non-targeting sgRNAs (10% of the library) were included as negative controls. These sgRNAs were cloned into a Bpu1102I/BstXI-linearized pBA904-mCherry plasmid backbone. This backbone was modified from pBA904 (Addgene, #122238) by replacing the BFP coding sequence with an mCherry fragment from pSLQ1651 (Addgene, #51024), using Gibson assembly. The library was amplified in *E. coli* (ensuring >500× coverage) and subjected to EndoFree maxiprep. The details of the library are provided in **Supplementary Table 7**.

#### Lentiviral transduction and CRISPRi screening

Lentiviral particles were produced in HEK293T cells co-transfected with the CRISPRi plasmid library, psPAX2, and pMD2.G using Lipofectamine 3000, and concentrated 100× with PEG8000. A total of 1×10⁷ K562-dCas9-GFP cells were lentivirally transduced with the CRISPRi sgRNA library at a MOI of ∼ 0.3 to ensure >500× sgRNA coverage. sgRNA-positive (GFP/mCherry-positive) cells were sorted by flow cytometry at Day 2–3 post-infection (defined as Day 0). For proliferation-based screening, cells were cultured for 28 days, and samples were collected on Day 0 and 28 for genomic DNA extraction and next-generation sequencing (NGS) to quantify sgRNA abundance. For FACS-based screening, CD71 (TFRC) surface expression was quantified by FACS at Day 14. Cells were stained with APC-conjugated anti-human CD71 antibody on ice for 15–20 min, washed, and sorted into CD71-high (top 30%) and CD71-low (bottom 30%) populations for genomic DNA extraction and NGS to determine sgRNA abundance. Two biological replicates were performed for these screens.

#### Data processing

Sequencing data generated from the TFRC-targeted CRISPRi screens (FASTQ format) were processed using MAGeCK to generate raw sgRNA read count tables. Raw log fold change (LFC) values for each sgRNA were calculated using MAGeCK. sgRNAs targeting the same CRE region were aggregated using the Robust Rank Aggregation (RRA) algorithm implemented in MAGeCK, and log10-transformed RRA scores (tRRA) were calculated to assess the statistical significance of each CRE in regulating cell proliferation or *TFRC* expression (**Supplementary Table 7**).

### CRISPR-Cas9-mediated enhancer deletion

#### sgRNA design and transfection

Paired sgRNAs flanking putative enhancer regions (designated as sgRNA-L and sgRNA-R) were designed using CHOPCHOP[92]. The pSpCas9(BB)-2A-GFP (PX458; Addgene #172221) and pSpCas9(BB)-2A-Puro (PX459; Addgene #48139) plasmids, each encoding either an L or R sgRNA for the corresponding candidate regulatory elements (CREs), were co-transfected into K562 cells using Lipofectamine 3000. Twenty-four hours post-transfection, the cells were selected with 2 μg/mL puromycin for 48 hours. For the enhancer region E37, which spans an extended sequence, an additional sgRNA (sgRNA-M) was designed to pair with sgRNA-L and sgRNA-R separately, thereby creating two distinct targeting combinations (L/M and M/R) to facilitate efficient deletion. For E37 targeting, cells were co-transfected with plasmid pairs expressing either sgRNA-L/M or sgRNA-M/R, followed by the same puromycin selection protocol. All sgRNA sequences are listed in **Supplementary Table 7**.

#### Single-cell cloning and genotyping

After puromycin selection, GFP+ cells were then sorted into 96-well plates by flow cytometry and expanded for ∼10 days. All genomic DNA was extracted from collected cells using the lysis buffer (300 mM NaCl, 0.2% SDS, 2 mM EDTA, 10 mM Tris-HCl pH 8.0) with 10 mg/mL RNase A, and DNA concentration was quantified using NanoDrop. Around 500 ng of genomic DNA were used to amplify the CRE deletion region using Phanta Super-Fidelity DNA Polymerase (Vazyme #P501-d3). The PCR primer sequences are listed in **Supplementary Table 7**.

### RT-qPCR

Total RNA was isolated from harvested cells using the UNlQ-10 Column TRIzol Total RNA Isolation Kit (Sangon Biotech, #B511321) according to the manufacturer’s instructions. For cDNA synthesis, 1 μg of total RNA was reverse-transcribed using random primers and MultiScribe reverse transcriptase (Thermo Fisher, #4311235) under the following conditions: 25 °C for 10 min, 37 °C for 90 min, and enzyme inactivation at 85 °C for 5 min. Quantitative PCR was performed using UltraSYBR 2× Master Mix (Sparkjade, #AH0105) on a QuantStudio 5 Real-Time PCR System (Thermo Fisher, #A28319). The thermal cycling protocol consisted of an initial denaturation at 95 °C for 10 min, followed by 40 amplification cycles of 95 °C for 10 s, 60 °C for 30 s, and 72 °C for 30 s. *RPS28* was used as the internal reference gene for normalization. All reactions were conducted in technical triplicates, and relative mRNA expression levels were calculated using the ΔΔCt method. The primers for qPCR are listed in **Supplementary Table 7**.

### Western blot

Cells were lysed in RIPA buffer (Beyotime, #P0013C) supplemented with a protease inhibitor cocktail (CWBIO, #CW2200S) and incubated on ice with gentle agitation for 20 min. Lysates were clarified by centrifugation at 14,000 × rpm for 10 min at 4 °C. The supernatants were collected, mixed with 5× SDS sample buffer, and denatured at 95 °C for 5 min. Samples were resolved by SDS–PAGE and transferred onto PVDF membranes. Membranes were probed with antibodies against CD71/transferrin receptor (ABclonal, #A22161), and α-tubulin (Proteintech, #11224-1-AP), followed by appropriate secondary antibodies. Signals were detected using a chemiluminescence substrate (Sparkjade, #ED0025) and captured with a Tanon imaging system (#5200).

### Cell proliferation assay

CRE knockout cells (5×10^3^ cells/well) and *TFRC* knockdown cells were seeded in triplicate in 96-well plates. At time points of Day 0, 1, 2, 3, and 4, add 10% volume of CCK8 reagent (MeilunBio #MA0218) and incubate for 1 h at 37°C. OD450 was measured to assess proliferation relative to Day 0.

### Iron uptake assay

FITC-conjugated holo-transferrin (Fe³⁺-Tf) was prepared by incubating 100 nM apo-transferrin with 200 nM ferric ammonium citrate (Sangon Biotech, #A500061) in serum-free medium at 37 °C for 1 h. Cells were first iron-depleted by treatment with 10 μM deferiprone (MedChemExpress, #HY-B0568) in serum-free medium for 12 h. The cells were then washed twice with PBS to remove residual deferiprone and incubated with 100 nM freshly prepared Fe³⁺-Tf in serum-free medium at 37 °C for 45 min to allow internalization. Following incubation, cells were washed twice with ice-cold PBS to remove surface-bound transferrin and immediately analyzed by flow cytometry. FITC fluorescence intensity was quantified from at least 10,000 single-cell events.

### shRNA mediated knockdown

shRNAs were designed using the GPP Web Portal (https://portals.broadinstitute.org/gpp/public/) and cloned into pLKO.1-puro vector (Addgene, #8453), individual shRNAs were lentivirally transduced into the cells at a high MOI (>1). Total RNA was extracted 2 days post-infection, and *TFRC* expression levels were quantified by RT–qPCR and cell proliferation was also measured. The sequences for shRNA are listed in **Supplementary Table 7**.

## Data and code availability

The data supporting the findings of this study are available within the paper and its Supplementary Information. The raw sequencing data generated in this study have been deposited in and are publicly accessible through the Genome Sequence Archive (GSA-Human) at the National Genomics Data Center, China National Center for Bioinformation / Beijing Institute of Genomics, Chinese Academy of Sciences, under the accession code: HRA016252. https://ngdc.cncb.ac.cn/gsa-human. All datasets for eScreen training were published previously (**Supplementary Table 1**). The code of eScreen was available via GitHub https://github.com/xmuhuanglab/eScreen. And user-friendly web server for online queries and visualization was available via https://escreen.huanglabxmu.com/.

## Authors’ Contributions

J.H., T.F., S.L., and L.L. conceived and designed the experiments. S.L., L.L., Z.L. and R.C. H.L. performed the computational analysis with help from Y.P. H.Z., Y.Y., Y.L., and Y.H. carried out experimental validation with help from X.N. F.C. and J.D. prepared the experimental materials and cultured the cell lines. S.L., L.L., J.H. and T.F. wrote the manuscript with other authors. J.H. and T.F. supervised the study.

## Acknowledgment

We thank Drs. Guo-Cheng Yuan and Jian Xu for useful comments and Mrs. Binbin Fan for technical support. We thank the Experimental Technology Center, College of Life and Health Sciences (Northeastern University), and Laboratory of Life and Medicine (Northeastern University) for instrumental and technical support. This work was supported by the National Natural Science Foundation of China (92474104, 32370586 to J.H.,), the Fundamental Research Funds for the Central Universities (20720230068 to J.H.) and the Wang Deyao Outstanding Graduate Scholarship Program of Xiamen University; the National Key Research and Development Program of China (2025YFA0921900 to T.F.) and the Construction Project of Liaoning Provincial Key Laboratory, China (2022JH13/10200026) to T.F.

---

**Extended Data Fig. 1.**
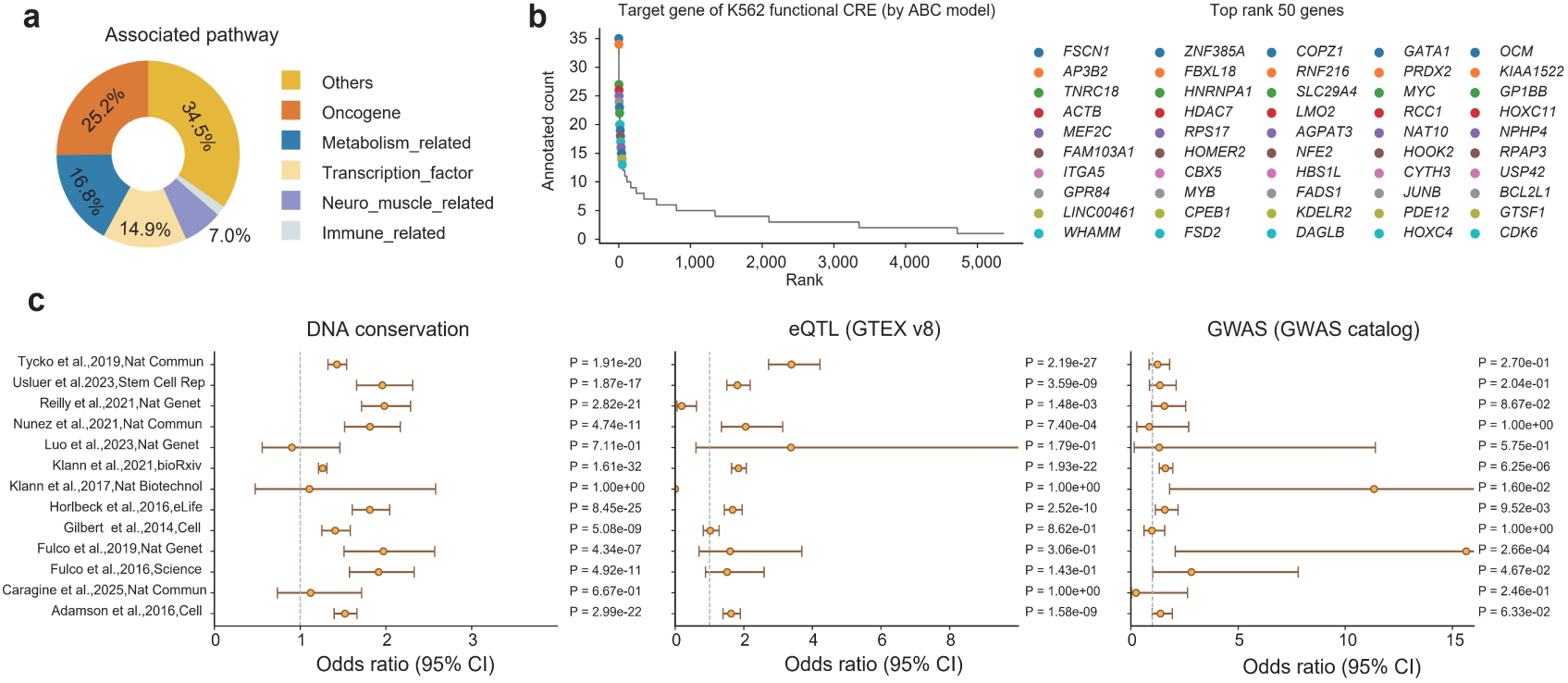
Characteristics of functional CREs identified across 379 CRISPR screens. **a**, Pie charts depict the distribution of functional CRE–associated genes across annotated biological pathways. **b**, Dot plot shows the distribution of associated genes of functional CREs in K562 cells. **c**, Enrichment of conserved elements, eQTLs, and GWAS loci among functional CREs identified from 13 K562 CRISPR screen datasets. Dots represent odds ratios (95% confidence intervals shown as bars), and significance was assessed using two-sided Fisher’s exact tests.

**Extended Data Fig. 2.**
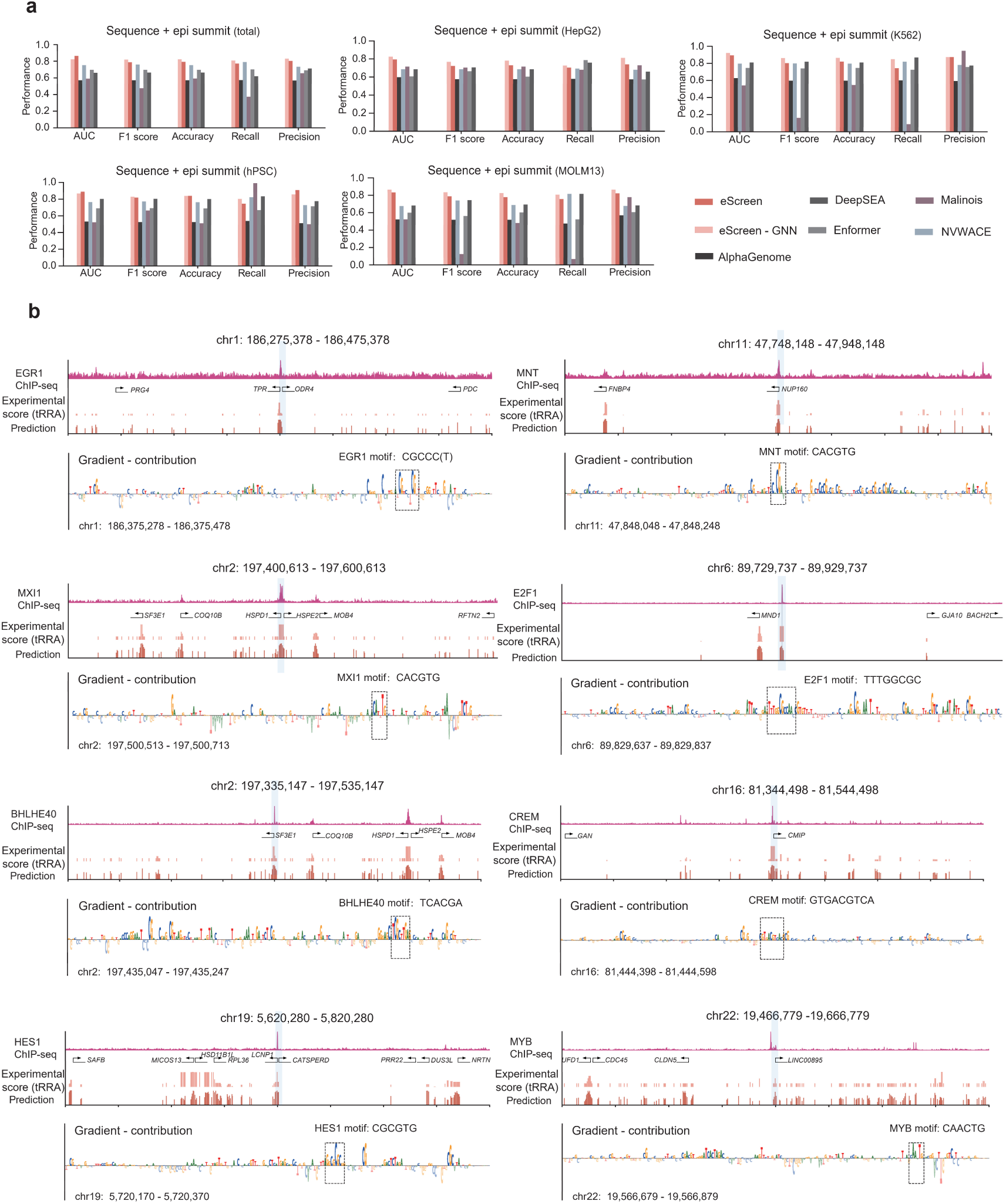
Benchmarking and mechanistic interpretation of eScreen predictions. **a**, Performance comparison of eScreen with eScreen-GNN, AlphaGenome, DeepSEA, Malinois, Enformer and NVWACE for the prediction of functional CREs. All models were trained and evaluated on functional CRE sequences across multiple cell lines. **b**, Representative examples of eScreen-predicted regulatory activity across diverse TF binding sites, shown with matched ChIP-seq profiles and CRISPR experimental scores (tRRA). Gradient contribution analysis pinpointed nucleotide-level determinants predictive of diverse TF binding at putative functional CREs, validated by independent ChIP-seq datasets.

**Extended Data Fig. 3.**
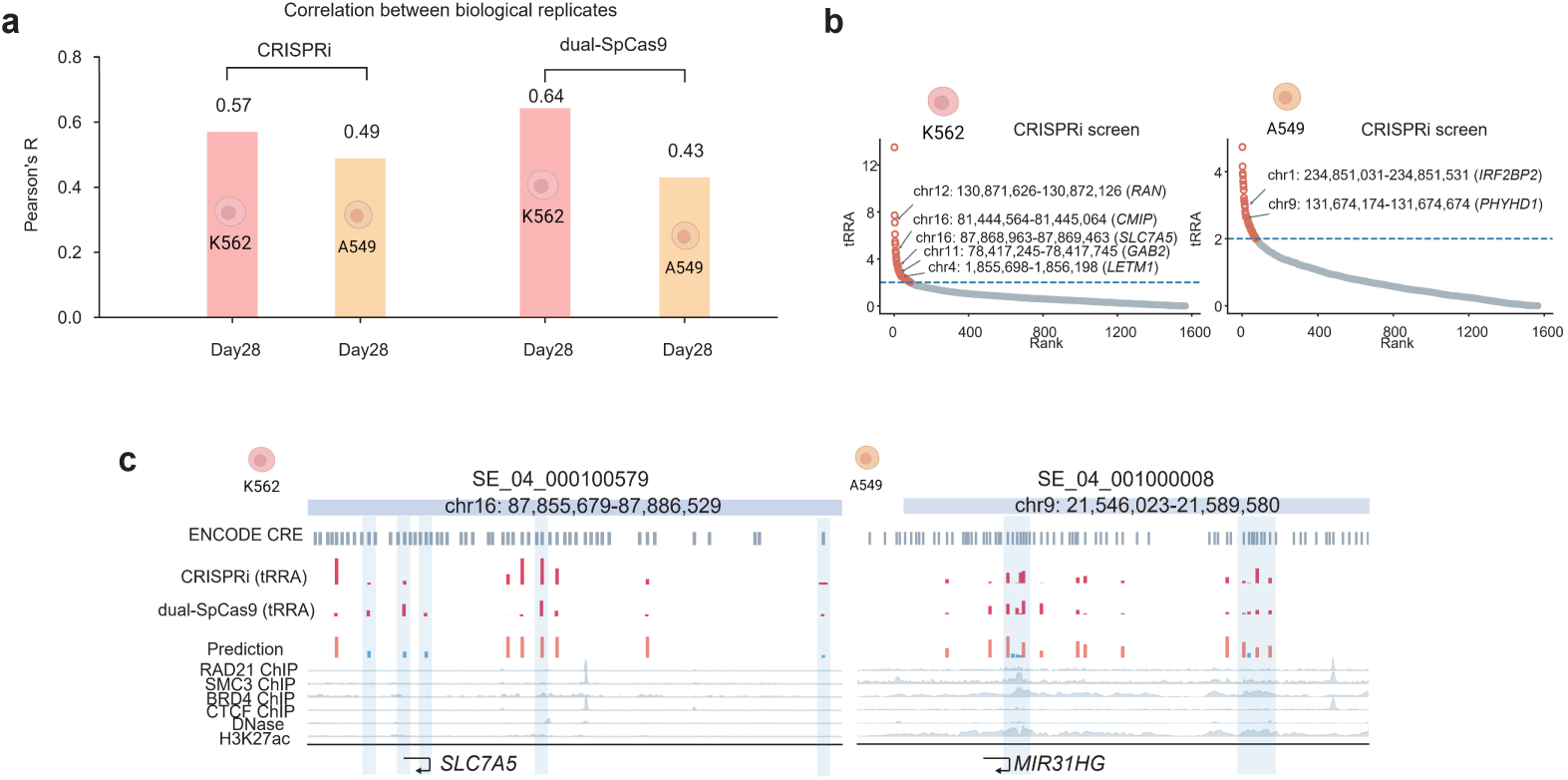
Benchmarking and mechanistic interpretation of eScreen predictions. **a**, Bar plot showing the correlation between biological replicates of CRISPRi and dual-SpCas9 screens in K562 and A549 cells. **b**, High-confidence functional CRE hits identified in K562 (left) and A549 (right) cells by CRISPRi screen. RRA plots display transformed RRA score (tRRA) from MAGeCK analysis (Day 28 vs. Day 0). The labeled genes are adjacent to CRE hits detected by the ABC model. **c**, Two representative super-enhancer loci showing eScreen-predicted CRE functional activity, experimental score (tRRA) across CRISPRi and dual-SpCas9, and matched epigenetic profiles for context.

**Extended Data Fig. 4.**
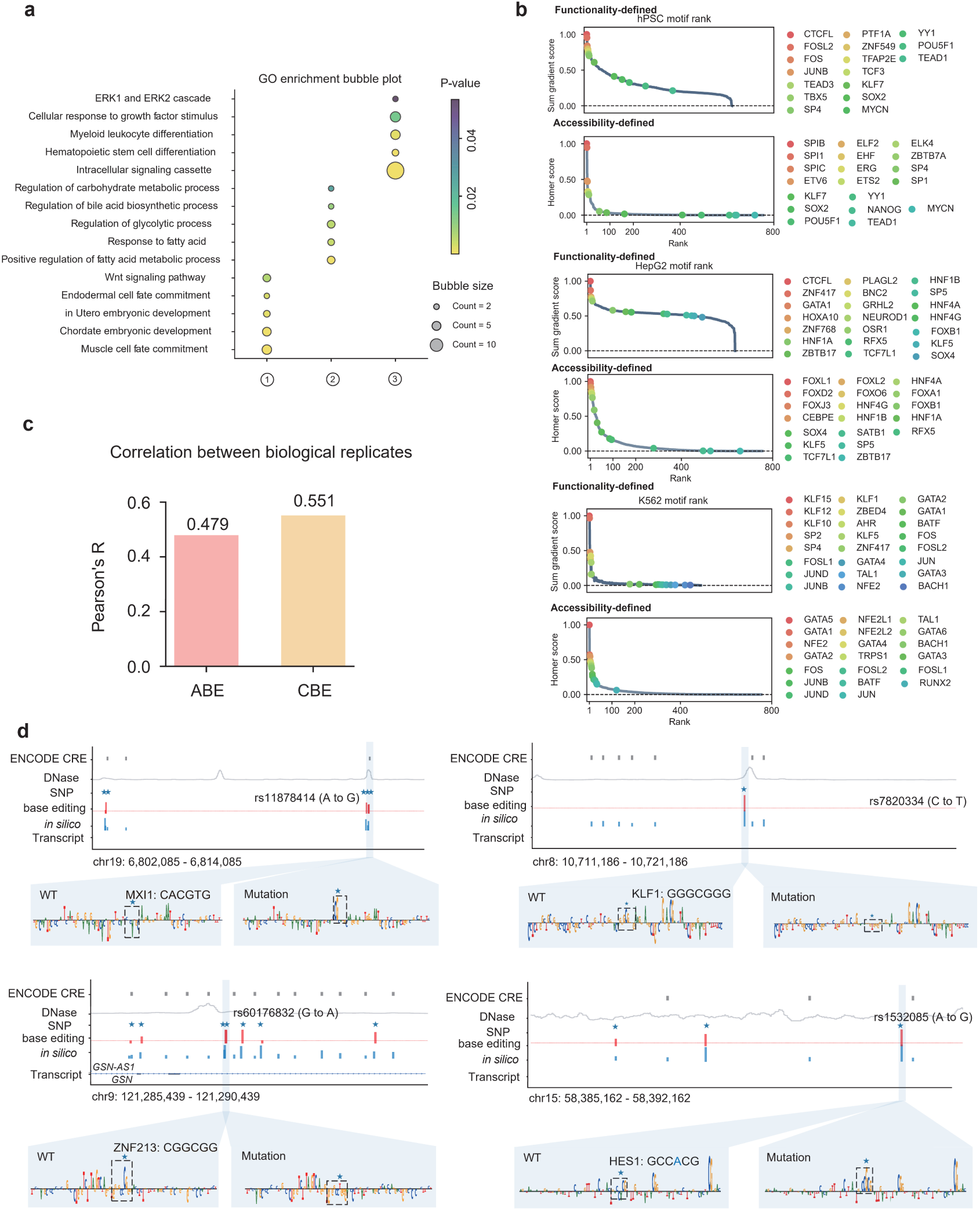
Functional contribution maps enhance detection of cell-type-specific regulatory motifs. **a**, Dot plot heatmap showing biological pathways significantly enriched for the distinct motif combination programs. Dot size indicates the number of associated motifs, and color represents the p-value. **b**, Dot plots comparing the ability of functionality-defined CREs (by eScreen) and accessibility-defined CREs (ATAC) to recover sequence features of cell-type-specific transcription factors across three CRE sources (hPSC, HepG2 and K562). **c**, Bar plot showing the correlation between biological replicates of ABE and CBE base editing screens in MOLM-13 cells. **d**, *In silico* nucleotide mutagenesis by eScreen of CREs surrounding rs11878414, rs60176832, rs7820334 and rs1532085, showing the predicted changes in regulatory activity (blue bar), and the corresponding phenotypic effect measured in base editing experiments (red bar). For CREs lacking SNPs, multiple A- or C- containing sequences were mutated, and the mean effect was quantified (blue bar without SNP).

**Extended Data Fig. 5.**
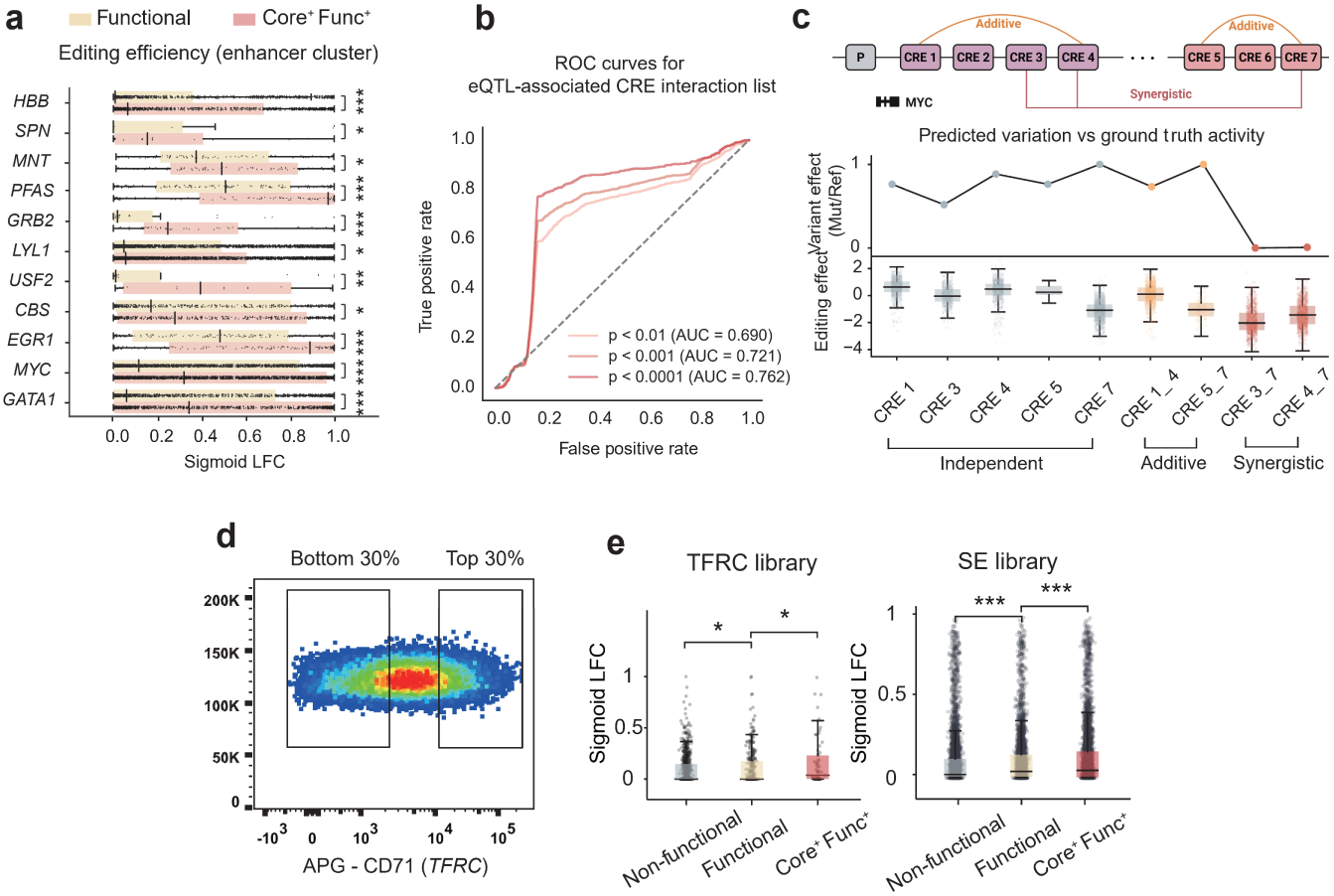
Validation of eScreen-predicted functional organization of enhancer cluster. **a**, Sigmoid LFC of core functional CREs and functional CREs in representative enhancer clusters. Significance was assessed using Mann-Whitney-U test (\**p* < 0.05, \*\**p* < 0.01, \*\*\**p* < 0.001; *n.*s., not significant). **b**, The ROC curve benchmarks *in silico* classified functional or non-functional CRE combinations against eQTL-inferred synergistic enhancer pairs used as the gold-standard reference set. **c**, *In silico* perturbation at the *MYC* locus reveals CRE synergistic interactions (e.g., E3-E7 or E4-E7), consistent with the patterns previously reported through multiplexed combinatorial CRISPR screening [69]. **d**, Density plot showing the flow cytometry gating strategy used to isolate CD71^high^ and CD71^low^ populations in the FACS-based CRISPRi screen. **e**, Comparison of editing efficiency (sigmoid LFC) among core functional, functional, and non-functional CREs in *TFRC* CRISPRi screen and SE screen (Fig. 3g). Significance was assessed using permutation tests (\**p* < 0.05, \*\**p* < 0.01, \*\*\**p* < 0.001; *n.*s., not significant).

**Extended Data Fig. 6.**
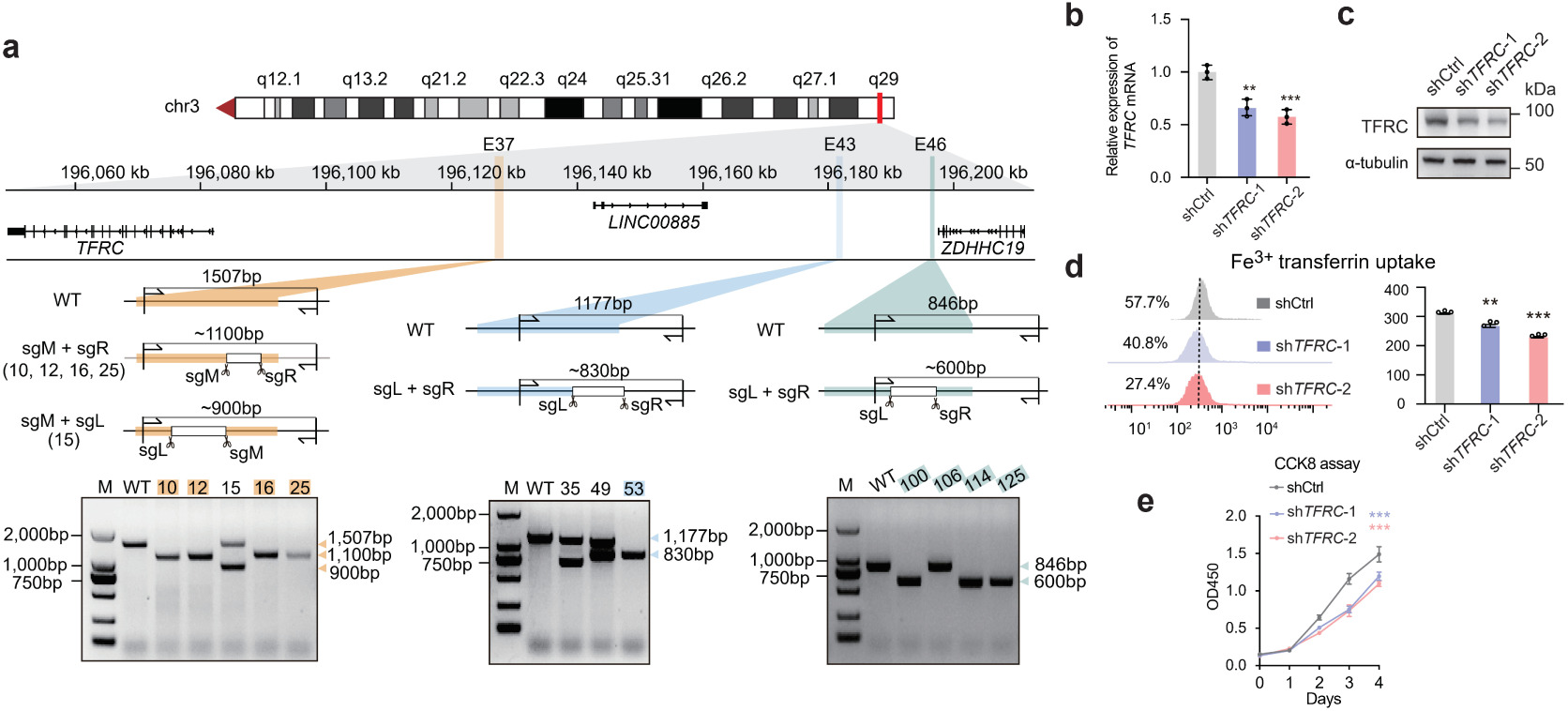
Knockout validation of regulatory activity within the *TFRC* enhancer cluster. **a-e**, Functional consequences of CRE knockout by SpCas9 and *TFRC* knockdown using shRNA. (**a**) Genotyping of single clonal CRE knockout cell lines. (**b-e**) Phenotypic assays following shRNA-mediated *TFRC* knockdown: (**b**) *TFRC* mRNA levels (RT-qPCR), (**c**) TFRC protein abundance (Western blot), (**d**) Cellular Fe^3+^ uptake, (**e**) Cell proliferation measured by CCK8 assay. For **b**-**e**, data were normalized to wild-type controls, and statistical significance was assessed by one-way ANOVA followed by Dunnett’s multiple comparisons test (\**p* < 0.05, \*\**p* < 0.01, \*\*\**p* < 0.001; *n.s.*, not significant).

